# Reciprocal-space mapping of diffuse scattering by serial femtosecond crystallography reveals analog-specific disorder in insulin analogs

**DOI:** 10.64898/2026.04.03.716400

**Authors:** Esra Ayan, Jungmin Kang, Takehiko Tosha, Makina Yabashi, Madan Kumar Shankar

## Abstract

Insulin detemir and insulin aspart are clinically complementary analogs engineered for distinct pharmacokinetic behavior, yet their comparative structural heterogeneity across temperature regimes remains insufficiently resolved. Here, we present a multi-scale crystallographic analysis integrating near-physiological serial femtosecond crystallography (SFX) with previously reported cryogenic and ambient multicrystal datasets for both analogs. Across conventional quality metrics, reciprocal-space intensity-field reconstructions, model-derived diffuse-scattering representations, Ramachandran stereochemical validation, solvent-accessibility coupling (*SA_Area_–MS_Area_*), and residue-level BDamage (a packing-normalized B-factor metric highlighting local mobility outliers) profiling, we identify a coherent ambient-versus-cryogenic contrast. Ambient datasets show broader reciprocal-space heterogeneity and more diffuse model-space distributions, consistent with increased conformational sampling outside cryogenic trapping. Despite this shared trend, disorder partitioning is analog-specific: detemir exhibits strong pseudo-translational signatures with moderate twinning, whereas aspart shows weak pseudo-translation but pronounced merohedral twinning approaching the theoretical twinned limit in ambient conditions. Importantly, backbone stereochemistry remains globally stable across all datasets, indicating that the observed differences reflect structured heterogeneity rather than model deterioration. Collectively, these findings support an ensemble-aware interpretation of insulin crystallography and provide transferable structural descriptors for analog comparison, stability assessment, and formulation-oriented design.

## Introduction

Insulin therapy remains central to diabetes management, and modern analog design has progressively separated basal and prandial functions through structure-guided molecular engineering. In this framework, long-acting analogs are optimized for delayed absorption and prolonged systemic residence, whereas rapid-acting analogs are optimized for accelerated dissociation and early post-injection activity. Among the most widely used examples, insulin detemir and insulin aspart represent distinct kinetic strategies that are clinically complementary rather than interchangeable. Detemir is acylated at LysB29 and formulated to support protracted action through self-association and reversible albumin interactions, while aspart (B28Pro→Asp) is engineered to weaken self-association and facilitate faster monomer availability after injection [1], [2], [3], [4], [5]. From a structural biology perspective, this pharmacological contrast is rooted in insulin’s polymorphic oligomerization landscape (monomer, dimer, and zinc-stabilized hexameric states with T/R-state variation), which is highly sensitive to pH, ligand environment, temperature, and crystal packing [6], [7].

Early crystallographic studies established the core architectures of both analogs, including detemir’s acylated framework and aspart’s hexameric assemblies in the trigonal setting [6], [8], [9]. These structural datasets have been indispensable for understanding analog behavior at atomic resolution, but they are dominated by cryogenic single-crystal measurements and therefore primarily report low-temperature, selected lattice states. Cryogenic crystallography is extremely powerful for high-resolution model building, yet it can attenuate thermally accessible conformational heterogeneity and may under-represent lattice-state diversity present near physiological conditions [10], [11]. By contrast, room-temperature and serial approaches can increase sensitivity to ensemble-level variability in both reciprocal-space sampling and disorder signatures. This distinction is especially relevant for insulin analogs, where comparatively subtle shifts in oligomerization equilibrium and local flexibility can influence macroscopic pharmacology. Accordingly, ambient-condition structural analysis is not intended to replace cryogenic structures, but to complement them by probing a broader functional conformational landscape [10], [11], [12], [13].

Bragg data provides information on the mean electron-density model, while diffuse intensity carries information on correlated deviations from that mean. These representations are informative for lattice sampling, anisotropy, and model–data consistency, but they are not equivalent to detector-level experimental diffuse-scattering measurements and should be interpreted with this distinction explicitly stated [14], [15]. For insulin analogs crystallizing in trigonal/rhombohedral settings, translational non-crystallographic symmetry (tNCS) and merohedral twinning can further complicate intensity statistics and refinement behavior. Modern likelihood-based crystallographic practice has established that these effects are diagnostically tractable and interpretable when explicitly modeled, rather than treated as simple noise [16], [17], [18], [19]. Therefore, comparative analysis of detemir and aspart across cryogenic and ambient datasets can be used to distinguish shared temperature-dependent trends from analog-specific disorder partitioning. Used conservatively, such metrics can provide an additional bridge between global reciprocal-space behavior and residue-level structural vulnerability, particularly when interpreted together with stereochemical validation and solvent-accessibility trends [20], [21]. This multi-scale strategy is well aligned with current IUCr standards [22], [23], emphasizing transparent model limitations and calibrated mechanistic inference.

The present study applies this integrative framework to near-physiological SFX datasets of insulin detemir and insulin aspart and compares them against representative cryogenic and ambient multicrystal PDB structures. By combining conventional crystallographic quality indicators, reciprocal-space mapping, model-derived diffuse representations, atom-level SA–MS analysis, and RABDAM descriptors, we evaluate whether the two analogs share a common ambient-versus-cryogenic disorder signature while preserving analog-specific fingerprints consistent with their opposing kinetic design goals. Beyond structural interpretation, this comparison aims to identify practical, quantifiable readouts that may be useful for formulation screening, stability assessment, and analog engineering in translational insulin development.

## 1. Results

### 1.1. Crystallographic quality assessment of near-physiological temperature SFX detemir

We processed ambient-temperature serial femtosecond crystallography (SFX) data for detemir insulin at resolutions of 22.58–2.85 Å (**Table 1, Table S1, Table S3, Figure S1**). The crystals are indexed in space group `R 3: H`, with unit-cell parameters `a = b = 81.58 Å` and `c = 81.33 Å`. The dataset was highly complete with 100 % overall completeness and a mean signal-to-noise ratio `(<I/σ>`) of 3.7. Consistent with ambient-temperature SFX collection, no ice-ring artifacts were detected. The isotropic Wilson B factor was 11.93 Å², and Matthews coefficient analysis assuming 50% solvent content suggested that the asymmetric unit can accommodate approximately 200 amino-acid residues. No significant anomalous signal was observed. Analysis of density statistics and packing identified a pronounced translational non-crystallographic symmetry (tNCS) component. Patterson analysis revealed a strong off-origin peak at 40.66 Å with a height of 55.35% relative to the origin peak. The corresponding `p = 3.205 × 10`⁻⁵ indicates that this feature is unlikely to occur by chance, supporting the presence of pseudo-translational symmetry in the lattice. In the presence of tNCS, the structure factor can be expressed as *F*(*hkl*) = *F*_1_(*hkl*) [1 + *e*^2*πih*⋅*t*^], which produces a systematic modulation of reflection intensities (*I* ∝ |*F*|^2^). This behavior is structurally informative but is expected to perturb intensity statistics and may contribute to moderately elevated refinement *R* values; however, our final refinement statistics (`Rwork = 0.25` and `Rfree = 0.28`) remain within the expected range for structures with pronounced tNCS. Additional tests, including the L-test and Wilson-moment analysis, also indicate partial twinning. The observed `<I²>/²` ratio was 1.719 (vs. 2.0 for untwinned data) and mean `|L|` was 0.422 (vs. 0.500 expected). Together, these metrics are consistent with a possible twofold merohedral twin law (`h, -h-k, -l`), with estimated twin fractions of ∼20–24% from Britton’s and maximum-likelihood approaches. Accordingly, the twin law `-h, -h-k, -l` was applied during refinement to improve map interpretability and limit model bias. Merging statistics for the 22.58–2.85 Å dataset further supported data usability (**Table 1**). Although the overall multiplicity was moderate (39.0), key quality indicators remained acceptable: mean intensity 626.2, overall mean `I/σ(I)` 3.7, `Rmerge` 0.209, `Rmeas` 0.296, and overall `CC1/2` 0.783. Importantly, in the highest-resolution shell (2.95–2.85 Å), with `CC1/2 = 0.665` and `I/σ(I) = 3.0`, supporting reliable downstream refinement (**Table S1**).

**Table 1.** Data collection and refinement statistics.

| Data collection | SFX detemir | SFX aspart |
| --- | --- | --- |
| Beamline | SACLA (BL2) | SACLA (BL2) |
| PDB ID | 25HL | 25HF |
| Data collection temperature | 298 K | 298 K |
| Resolution (Å) | 19.54 - 2.85<br>(2.952 - 2.85) | 23.12 - 2.2<br>(2.279 - 2.2) |
| Space group | R3:H | R3:H |
| Unit cell | 81.58 81.58 81.33 90 90 120 | 80.08 80.08 38.36 90 90 120 |
| Multiplicity | 39 (27.4) | 50 (30.2) |
| Rsplit (%) | 64 (73.21) | 37.3 (40.4) |
| Completeness % | 100 (100.00) | 100 (100.00) |
| Mean I/sigma(I) | 3.4 (3.0) | 3.5 (1.8) |
| Wilson *B*-factor | 11.93 | 32.99 |
| CC1/2 | 0.78 (0.67) | 0.92 (0.71) |
| CC* | 0.94 (0.90) | 0.98 (0.91) |
| R-work | 0.25 (0.28) | 0.25 (0.39) |
| R-free | 0.28 (0.32) | 0.28 (0.45) |
| Total reflections | 413176 (14450) | 479800 (14093) |
| Unique reflections | 10503 (528) | 9572 (466) |
| Protein residues | 200 | 100 |
| RMS (bonds) | 0.004 | 0.003 |
| RMS (angles) | 0.71 | 0.61 |
| Ramachandran favored (%) | 98.37 | 98.90 |
| Ramachandran allowed (%) | 0.54 | 0.56 |
| Ramachandran outliers (%) | 1.09 | 0.54 |
| Average *B*-factor | 34.6 | 31.27 |
| solvent | 33.19 | 34.05 |

**CC\*** is a metric derived from CC1/2 that estimates how well the measured diffraction data correlate with the true underlying signal.

### 1.2. Comparison of all detemir crystal structures along the three-fold symmetry axis

When the present ambient-temperature SFX dataset is compared with previously reported detemir structures (PDB IDs: 1XDA, 8HGZ, 9LVC, and 9LVX; **Figure S1**), notable differences appear in both lattice dimensions and diffraction statistics. Cryogenic single-crystal datasets (8HGZ, 9LVC) show relatively compact unit cells (`a = b = 78.885 Å` and `79.064 Å`), whereas the room-temperature multicrystal dataset 9LVX is expanded (`a = b = 80.898 Å`), and the current SFX data is larger still (`a = b = 81.58 Å`). This trend is consistent with lattice relaxation at ambient temperature. Despite this expansion, tNCS is consistently observed in detemir crystals in space group `R3`. Strong off-origin Patterson peaks are present in all datasets, including cryogenic structures (59.23% in 8HGZ; 58.98% in 9LVC) and room-temperature datasets (75.39% in 9LVX; 55.35% in the present SFX data), indicating that pseudo-translation is likely an intrinsic packing feature.

The clearest contrast is in twinning behavior. Cryogenic single-crystal datasets (8HGZ, 9LVC) display near-untwinned intensity statistics (mean `|L| = 0.504` and `0.511`), whereas 9LVX shows marked deviation `(|L| = 0.407`) and reported merohedral twinning with operator (`h, −h−k, −l`) and twin fraction up to ∼37.6%. The present SFX dataset follows the same pattern `(|L| = 0.422`), supporting a similar twin operator with an estimated twin fraction of ∼20–24%. Overall, these findings suggest that detemir crystals may be more susceptible to merohedral twinning under ambient-temperature and/or multicrystal data-collection conditions. Such behavior is well documented for trigonal and rhombohedral crystal systems, in which lattice symmetry can permit overlap of symmetry-related twin domains [24], [25]. Because conventional cryogenic single-crystal workflows often prioritize the most well-ordered crystals, these defects may be systematically underrepresented. By contrast, SFX and related room-temperature multicrystal approaches sample broader crystal populations and therefore offer a more comprehensive view of intrinsic lattice heterogeneity, including the underlying propensity for twinning.

### 1.3 Crystallographic quality assessment of near-physiological temperature SFX aspart

The near-physiological-temperature SFX dataset for aspart was processed to 2.20 Å (23.12–2.20 Å) (**Table 1, Table S2, Table S3, Figure S2**) and indexed in the trigonal space group `R3:H` (No. 146), with unit-cell parameters `a = b = 80.08 Å` and `c = 38.36 Å`. Data completeness is essentially 100% across the full resolution range. The overall mean signal-to-noise ratio`, I/`σ(I)⟩ ≈ 3.5, indicates reliable data quality under ambient-temperature SFX conditions. Merging statistics are likewise consistent with a high-quality dataset, with mean intensity ∼1181.0`, I/σ(I) = 3.5, Rmerge = 0.151, Rmeas = 0.214,` and `CC1/2 = 0.916`. In the highest-resolution shell (2.28–2.20 Å), the data remain informative `(CC1/2 = 0.712,` ⟨`I/σ(I)`⟩ `= 1.8),` supporting retention of the full `2.20 Å` cutoff for refinement. Wilson analysis yielded an isotropic *B*-factor of `32.20 Å²,` consistent with moderate atomic displacement parameters expected for room-temperature crystallography. Matthews coefficient analysis, assuming approximately `50%` solvent content, indicates that the asymmetric unit contains about 87 amino-acid residues, in agreement with the expected molecular composition. Electron-density statistics and crystal-packing analysis show no evidence of translational non-crystallographic symmetry (tNCS). Patterson analysis yields the strongest off-origin peak at `∼7.7%` of the origin peak, at a distance of `16.81 Å (p ≈ 0.38)`, indicating that pseudo-translation is not significant in this lattice.

By contrast, intensity-distribution analyses indicate pronounced merohedral twinning. The observed ratio ⟨`I²`⟩`/`⟨`I`⟩`² = 1.527` deviates substantially from the untwinned expectation (2.0) and approaches the theoretical limit for perfect twinning (1.5). Additional moment-based indicators support this interpretation. Collectively, these results are consistent with strong twinning, likely associated with the twofold merohedral twin operator (`h,−h−k,−l`) and a twin fraction approaching `0.5.` This twin law was therefore included during refinement to account for the twinned contribution and reduce model bias [25]. Overall, despite strong merohedral twinning, the dataset provides high completeness, stable signal-to-noise, and meaningful high-resolution information, supporting robust structure determination [25], [26], [27] at near-physiological temperatures.

### 1.4. Comparison of all aspart crystal structures along the three-fold symmetry axis

Comparison of the ambient-temperature SFX aspart dataset with previously reported structures (4GBK, 4GBC, and 8Z4B), all indexed in space group `R3:H` (No. 146), indicates systematic differences in lattice dimensions, diffraction statistics, and twinning behavior, despite broad overall crystallographic similarity. The unit-cell parameters show a clear trend. The cryogenic single-crystal structures 4GBK (`a = b = 77.50 Å`) and 4GBC (`a = b = 78.15 Å`) are relatively compact, whereas 8Z4B (`a = b = 80.37 Å`) and the ambient-temperature SFX dataset (`a = b = 80.08 Å`) show expanded lattices. This increase is consistent with lattice relaxation under non-cryogenic, near-physiological conditions, as commonly observed in room-temperature crystallography. Despite this expansion, all datasets show no substantial evidence for translational non-crystallographic symmetry (tNCS). Patterson analyses report weak off-origin peaks with relative heights of approximately `7.20%` (4GBK), `8.04%` (4GBC), `8.83%` (8Z4B), and `7.69%` (SFX), each associated with a high `p` value. These observations suggest that pseudo-translation is not a dominant feature of aspart crystal packing, in contrast to some related insulin-analog systems. This behavior is compatible with the distinct packing arrangement of aspart insulin in the *T_3_R_3_* conformational state, which favors rapid monomer–hexamer exchange and may limit the emergence of strongly correlated pseudo-translational relationships [6].

The most marked difference among the datasets is observed in twinning behavior. The single-crystal cryogenic structures 4GBK and 4GBC show near-untwinned intensity statistics, with ⟨`I²`⟩`/`⟨`I`⟩`²` values of `1.824` and `1.771`, respectively, consistent with expected `L-test` characteristics for predominantly single-domain crystals. By comparison, 8Z4B (⟨`I²`⟩`/`⟨`I`⟩`² = 1.522`) and the SFX dataset (`1.527`) both approach the theoretical value of `1.5` for perfectly twinned crystals, indicating pronounced merohedral twinning. In both cases, the twinning is consistent with the twofold merohedral operator (`h,−h−k,−l`), with estimated twin fractions near `0.5`. These findings suggest that aspart crystals (similarly, detemir crystals in rhombohedral packing) formed under ambient or near-physiological conditions may be intrinsically more susceptible to merohedral twinning [25], [27]. A plausible interpretation is that increased conformational flexibility of the aspart molecule, together with greater lattice plasticity in the *T_3_R_3_* state, facilitates symmetry-related domain intergrowth during crystal growth. Conversely, conventional cryogenic single-crystal workflows, which often include manual selection of morphologically well-ordered crystals, may under-sample such lattice imperfections.

Taken together, these results indicate that SFX and room-temperature approaches provide a broader and potentially more representative view of the intrinsic crystallographic heterogeneity of aspart insulin. In particular, they reveal a pronounced propensity for merohedral twinning that appears less evident in conventional cryogenic crystallographic datasets.

### 1.5. Reciprocal space mapping of lattice heterogeneity in three-fold symmetric crystals

Reciprocal-space representations generated from observed Bragg intensities and model-derived calculated structure factors show clear differences among the detemir datasets collected under different temperature and sampling conditions (**Fig. 1**) [28], [29], [30]. In the three-dimensional reciprocal-lattice point clouds, the ambient-temperature SFX (**Fig. 1A**) and multicrystal 9LVX (**Fig. 1C**) datasets appear sparser and more heterogeneous than the cryogenic single-crystal datasets 8HGZ (**Fig. 1B**) and 9LVC (**Fig. 1D**), indicating differences in reciprocal-space sampling and reflection distributions [29], [30]. These differences are consistent with the resolution limits of the datasets (*q* = 1/*d*): the cryogenic structures 8HGZ (1.70 Å) and 9LVC (2.00 Å) extend to higher reciprocal-space frequencies (*q* ≈ 0.6 Å^−1^), whereas the ambient-temperature SFX (2.85 Å) and 9LVX (2.37 Å) datasets are restricted to lower *q* values (*q* ≈ 0.35 − 0.40 Å^−1^), resulting in a visibly more confined reciprocal-space volume. Consistent with this observation, the *HK*0 and 0*KL* section maps indicate that the ambient-temperature datasets (**Fig. 1A_i-ii_** and **Fig. 1C_iii-iv_**) display less uniform and more heterogeneous reciprocal-space intensity patterns than their cryogenic counterparts (**Fig. 1B_i-ii_** and **Fig. 1D_iii-iv_**) [28], [29]. Since these maps were reconstructed from merged Bragg-reflection data using Gaussian-kernel broadening, rather than from raw detector diffuse-scattering images, they should be interpreted as regularized reciprocal-space representations and not as direct measurements of diffuse scattering. In macromolecular crystallography, reciprocal-space mapping remains informative for visualizing lattice variation, reflection broadening, and reciprocal-space heterogeneity [28], [29]. Within this framework, the greater heterogeneity observed in the ambient-temperature datasets is consistent with independent crystallographic evidence for increased lattice disorder under non-cryogenic and multicrystal conditions [30]. Furthermore, the lattice expansion observed at ambient temperature (*e.g.,* `a = b = 81.58 Å`) leads, by the inverse relationship between real and reciprocal space, to a closer spacing of Bragg reflections within the confined reciprocal-space volume. In addition, the periodic alternation of stronger and weaker layers visible in the 0*KL* projections [31] is compatible with previously identified pseudo-translational modulation in rhombohedral detemir crystals. Taken together, the comparative reciprocal-space analysis indicates that ambient-temperature multicrystal approaches, including SFX, capture a broader range of crystal-to-crystal variability than conventional cryogenic single-crystal measurements [30] [32].

**Figure 1.**
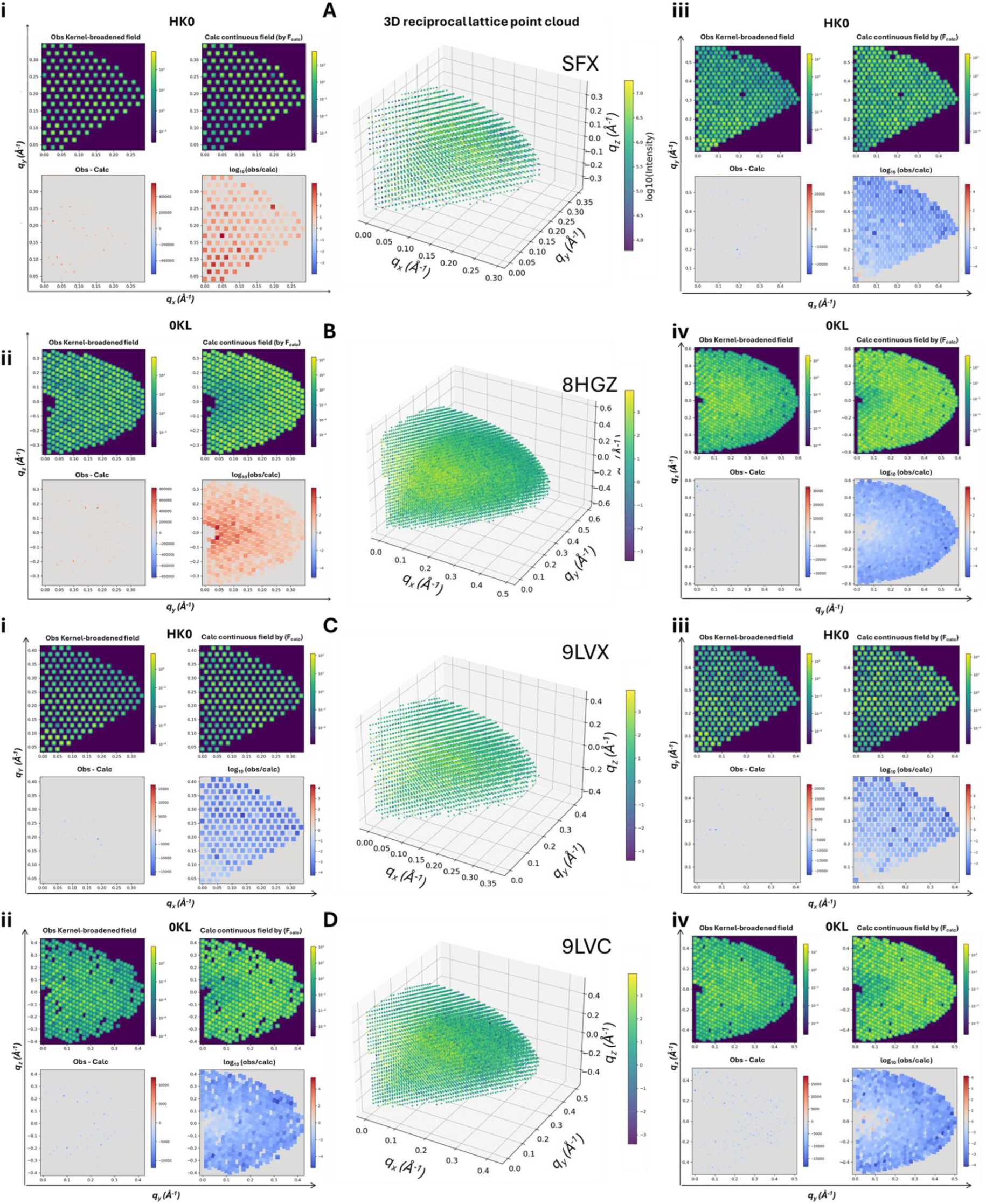
Comparative reciprocal-space representations of detemir insulin datasets collected under different temperature and sampling conditions. Three-dimensional reciprocal-lattice point clouds and two-dimensional sections (insets i-iv for *HK0* and *0KL*) are shown for four datasets: **(A)** ambient-temperature SFX, **(B)** cryogenic single-crystal 8HGZ, **(C)** ambient-temperature multicrystal 9LVX, and **(D)** cryogenic single-crystal 9LVC. For each dataset, observed Bragg intensities are displayed as Gaussian-kernel-broadened fields (*Obs*), alongside model-derived continuous intensity distributions from structure factors (*F_calc_*), with corresponding difference maps (*Obs − Calc*) and log-ratio maps [*log_10_(Obs/Calc)*]. The three-dimensional point clouds show dataset-dependent reciprocal-space coverage consistent with resolution limits: cryogenic datasets extend to larger reciprocal-space vectors (*q*), whereas ambient-temperature datasets are confined to lower *q* values and appear sparser and more heterogeneous. The *HK0* and *0KL* sections further illustrate differences in intensity distribution and uniformity. As these maps are reconstructed from merged Bragg reflections using kernel broadening, they are interpreted as regularized reciprocal-space intensity representations rather than direct diffuse-scattering measurements. The periodic modulation in the *0KL* sections is consistent with pseudo-translational symmetry in rhombohedral detemir crystal packing.

Likewise, to assess the diffraction characteristics of aspart insulin across different temperature and data-collection conditions, reciprocal-space representations were generated for the ambient-temperature SFX dataset and compared with previously reported structures [4GBK and 4GBC (cryogenic single crystal), and 8Z4B (ambient temperature)]. Three-dimensional reciprocal-lattice point clouds, together with HK0 and 0KL sections, were analyzed (**Fig. 2**). The three-dimensional distributions show differences in reciprocal-space coverage that are consistent with the respective resolution limits (**Fig. 2A**). The highest-resolution cryogenic dataset, 4GBC (1.78 Å), extends to the largest reciprocal-space vectors (*q*) (**Fig. 2C**), whereas the ambient-temperature SFX dataset (2.20 Å) extends further than the cryogenic 4GBK dataset (2.40 Å) (**Fig. 2B**). Accordingly, in contrast to the detemir comparison, reciprocal-space extension in aspart is not separated strictly by temperature but is more closely related to the intrinsic diffraction limit of each crystal. More pronounced differences are observed in intensity distributions. In the cryogenic single-crystal datasets (4GBK and 4GBC), Bragg reflections form well-defined and regularly spaced intensity maxima, consistent with comparatively uniform reciprocal-space sampling (**Fig. 2B-C**). By contrast, the ambient-temperature datasets (SFX and 8Z4B) show substantially greater heterogeneity (**Fig. 2A** and **Fig. 2D**). This is also evident in the corresponding difference representations [`Obs − Calc` and `log10(Obs/Calc)`], which show a broader spread of deviations for the ambient-temperature datasets. This increased heterogeneity is consistent with strong merohedral twinning. Whereas 4GBK and 4GBC are effectively untwinned, both SFX and 8Z4B show substantial twin fractions (up to approximately 49%) under the twin operator (`h,−h−k,−l`), which can lead to reciprocal-lattice overlap and perturbation of regular intensity distributions. The comparative analysis indicates that ambient-temperature multicrystal approaches, including SFX, sample a broader and more heterogeneous reciprocal-space distribution [9], [30] consistent with the intrinsic structural plasticity and twinning propensity [26] of aspart insulin [6] under near-physiological conditions.

**Figure 2.**
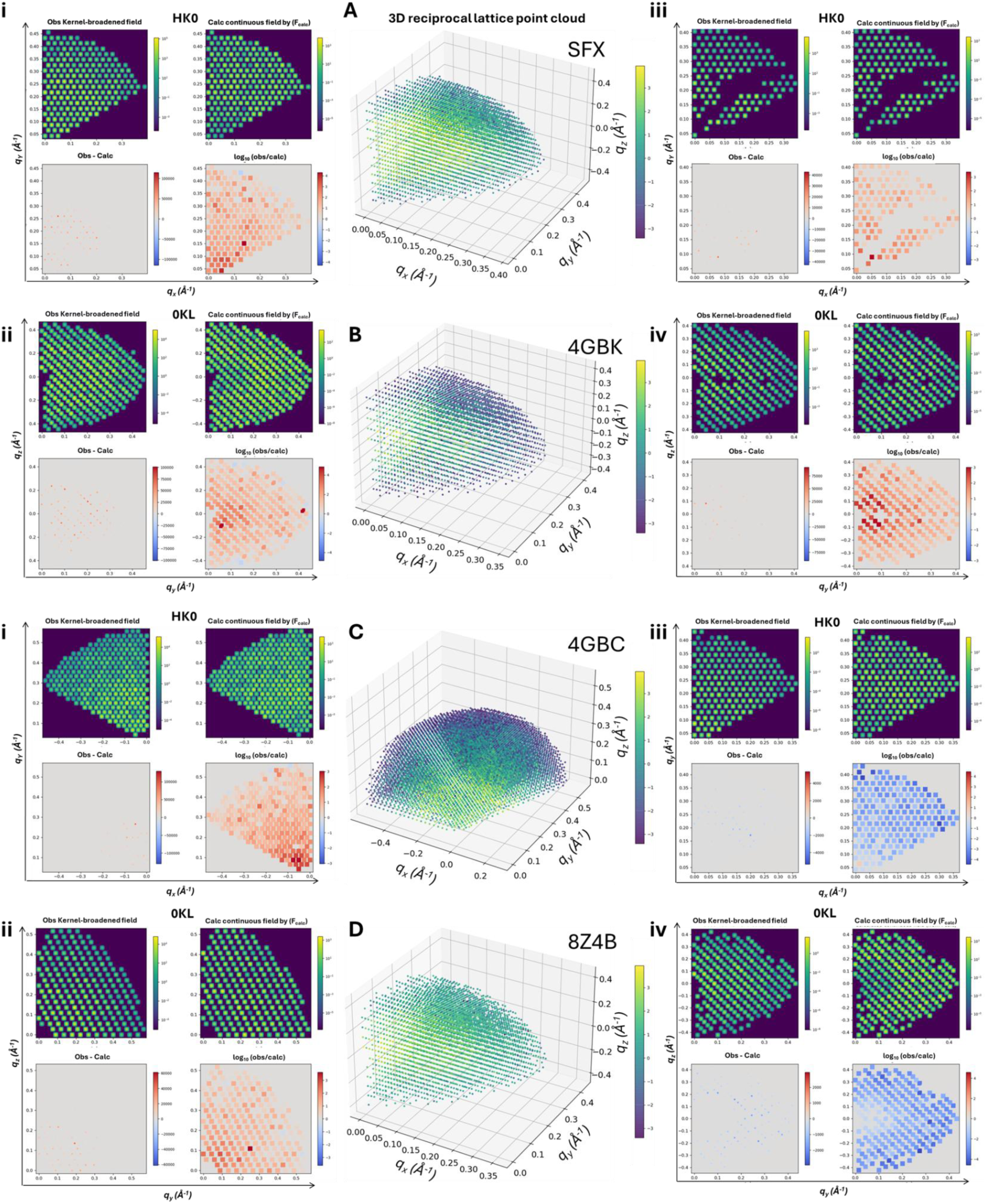
Comparative reciprocal-space representations of aspart insulin datasets collected under different temperature and sampling conditions. Panels A_i–ii_: ambient-temperature SFX (2.20 Å), with HK0 (i), three-dimensional point cloud (A), and 0KL (ii). Panels B_iii-iv_: cryogenic single-crystal 4GBK (2.40 Å). Panels C_i-ii_: cryogenic single-crystal 4GBC (1.78 Å). Panels D_iii-iv_: ambient-temperature multicrystal 8Z4B (2.30 Å). HK0 and 0KL sections are shown as Observed (*Obs*) Gaussian-kernel-broadened fields, Calculated (*Calc*) continuous fields, linear difference maps (*Obs − Calc*), and logarithmic ratio maps [*log_10_(Obs/Calc)*]. The reciprocal-space volume in the three-dimensional point clouds (A, B, C, D) follows dataset-specific diffraction limits (*q* range). In aspart, reciprocal-space extent is therefore governed primarily by crystal diffraction limit rather than temperature alone: SFX (2.20 Å) extends beyond 4GBK (2.40 Å), whereas the largest extent is observed for 4GBC (1.78 Å). Section maps (HK0 and 0KL), especially Obs − Calc maps, indicate clear differences in reciprocal-space homogeneity. These two-dimensional sections are regularized maps reconstructed from merged Bragg reflections using Gaussian kernel broadening, and do not represent direct measurements of raw diffuse scattering.

Collectively, comparative reciprocal-space mapping of detemir and aspart insulin supports the utility of SFX and multicrystal approaches at near-physiological temperature for resolving lattice heterogeneity [28]. Compared with conventional cryogenic single-crystal datasets, both analogs exhibit greater reciprocal-space heterogeneity and a sparser reflection distribution at ambient temperature. The crystallographic character of this disorder, however, might appear to be molecule-dependent. In detemir (**Fig. 1**), the reciprocal-space heterogeneity is consistent with periodic intensity modulation associated with pseudo-translational symmetry (tNCS), together with previously identified merohedral twinning [8]. In aspart (**Fig. 2**), the broader and less regular intensity variation is consistent with strong merohedral twinning, as supported by independent crystallographic analyses [26]. Given the analytical value of reciprocal-space mapping in protein crystallography for visualizing lattice defects and reflection broadening [28], this integrated analysis indicates that insulin analogs at near-physiological temperature exhibit not only volumetric lattice relaxation but also an intrinsic propensity for twinning, potentially linked to molecular flexibility. Furthermore, the results support the view that ambient-temperature multicrystal approaches sample lattice variation more comprehensively than conventional cryogenic single-crystal experiments [6], [9].

### 1.6. Ramachandran analysis of structural plasticity and local backbone integrity

To evaluate local backbone geometry and stereochemical quality in detemir insulin models determined under different temperature and data-collection conditions (ambient-temperature SFX and 9LVX; cryogenic 8HGZ and 9LVC), Ramachandran plots were analyzed (**Fig. 3, Fig S3**). Separate contour and scatter plots for general residues, glycine, and proline show that, across all datasets, main-chain ϕ and ψ torsion angles are concentrated in energetically favorable regions and remain within accepted stereochemical limits [33]. In particular, glycine residues, which have greater conformational freedom, and proline residues (*n* = 4), which are conformationally constrained, both agree well with residue-specific Ramachandran distributions, without clear outliers beyond classical allowed boundaries [33], [34]. Comparison across datasets shows the expected resolution-dependent trend. The cryogenic single-crystal datasets (8HGZ and 9LVC; **Fig. 3B, D**), which are higher-resolution, display relatively tighter clustering in ϕ/ψ space [34]. By contrast, ambient-temperature multicrystal datasets (SFX and 9LVX; **Fig. 3A, C**) show relatively broader angular distributions, consistent with increased thermal motion. Nevertheless, residues do not populate clearly disallowed regions in any dataset. These observations indicate that, despite pronounced lattice heterogeneity and twinning detected in reciprocal-space analyses, the local three-dimensional fold of detemir remains stereochemically stable under both temperature regimes.

**Figure 3.**
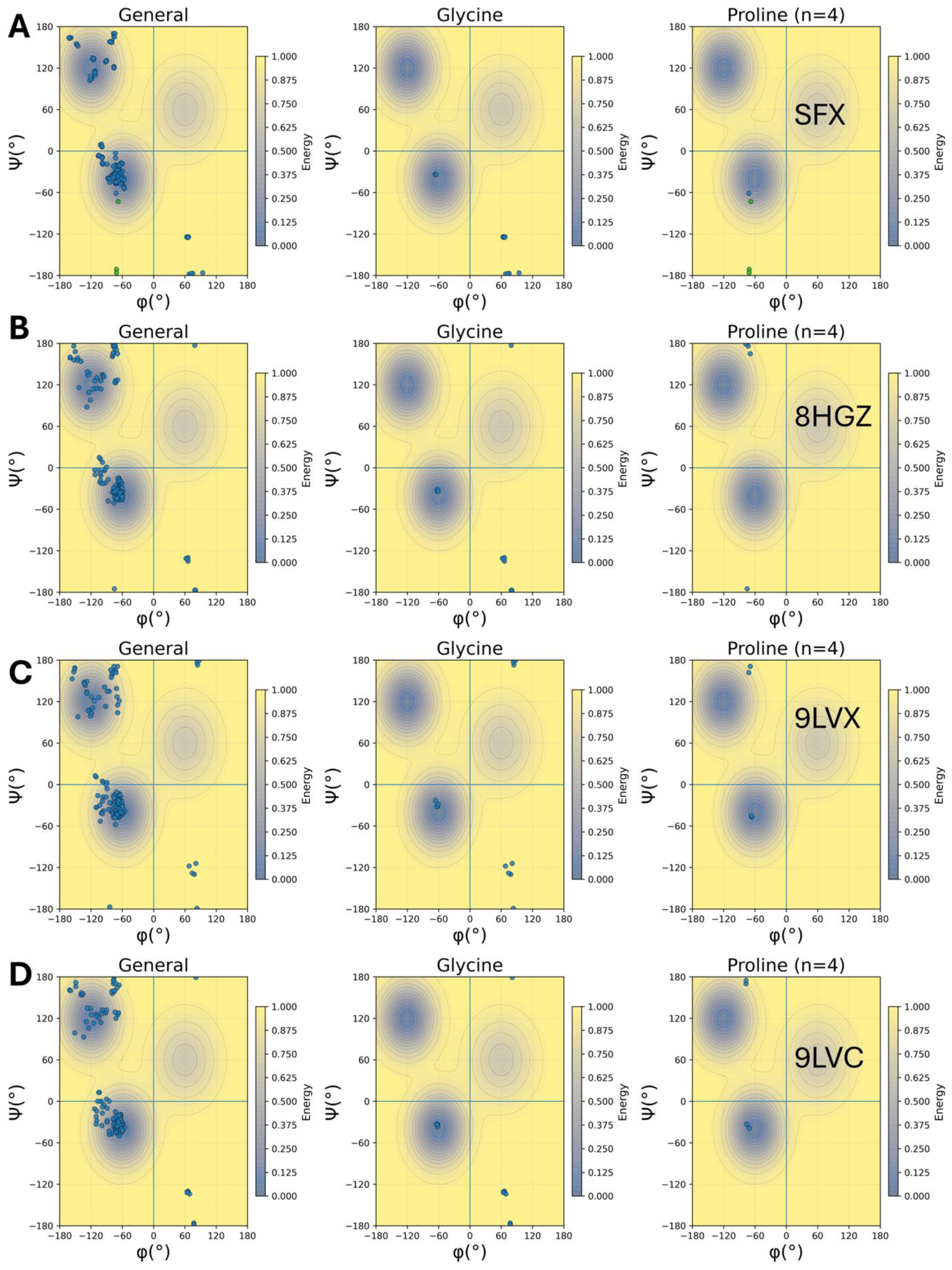
Ramachandran analysis of detemir insulin structures determined under different temperature and data-collection conditions. (A) Ambient-temperature serial femtosecond crystallography (SFX, 2.85 Å), (B) cryogenic single-crystal dataset (8HGZ, 1.70 Å), (C) ambient-temperature multicrystal dataset (9LVX, 2.37 Å), and (D) cryogenic single-crystal dataset (9LVC, 2.00 Å), showing distributions of ϕ and ψ torsion angles. In each panel, residues are shown separately as general amino acids (General), glycine (Glycine), and proline [Proline, n = 4]. Background contour maps indicate energetically favored and allowed regions of Ramachandran space, and observed ϕ/ψ values are shown as scatter points. Across all datasets, the large majority of residues occupy favored regions, and expected residue-specific distribution patterns are retained.

Similarly, to evaluate local backbone geometry and stereochemical quality in aspart insulin models determined under different temperature and data-collection conditions (ambient-temperature SFX and 8Z4B; cryogenic single-crystal 4GBK and 4GBC), Ramachandran plots were also analyzed (**Fig. 4, Fig. S4**). Contour maps generated separately for general residues, glycine, and proline indicate that, in all datasets, main-chain ϕ and ψ angles are concentrated in energetically favored regions and remain within accepted steric limits [33]. Unlike detemir, aspart insulin carries a B28 Pro→Asp substitution, and therefore contains no proline at that position; accordingly, no residues are present in the proline-specific distribution in these models (*n* = 0) [35]. Glycine residues (*n* = 6) agree with the broader glycine-specific Ramachandran distribution and remain within classical allowed limits [34]. Across datasets, a resolution- and condition-dependent trend is apparent. The highest-resolution cryogenic dataset, 4GBC (1.78 Å), shows the relatively tightest clustering in ϕ/ψ space (**Fig. 4C**) [34], followed by lower-resolution cryogenic dataset 4GBK (2.40 Å) (**Fig. 4B**). By contrast, broader angular distributions are observed in ambient-temperature multicrystal datasets [SFX (2.20 Å) and 8Z4B (2.30 Å)] (**Fig. 4A, D**). Nevertheless, no clear outliers are observed in disallowed regions in any dataset. These results indicate that, despite differences in resolution and data-collection regime, the local three-dimensional fold of aspart insulin remains stereochemically stable under both ambient and cryogenic conditions.

**Figure 4.**
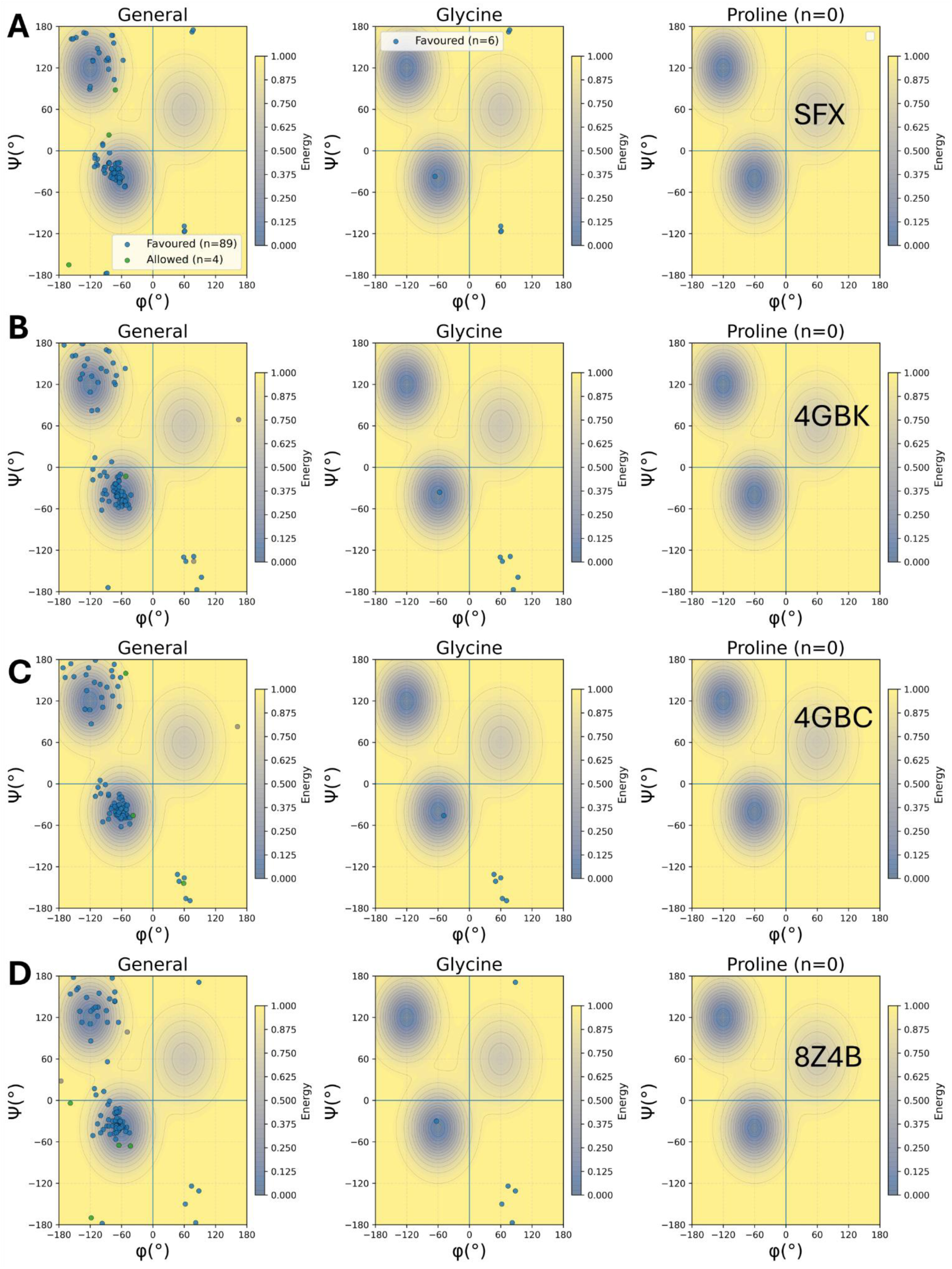
Ramachandran analysis of aspart insulin models determined under different data-collection conditions: (A) ambient-temperature SFX (2.20 Å), (B) cryogenic 4GBK (2.40 Å), (C) cryogenic 4GBC (1.78 Å), and (D) ambient-temperature multicrystal 8Z4B (2.30 Å). Distributions of ϕ/ψ angles are shown for general residues and glycine; proline plots are empty in all aspart datasets (n = 0) because of the B28 Pro→Asp substitution. Across datasets, residues are concentrated in favored/allowed regions, with tighter clustering in high-resolution 4GBC and broader spread in SFX, 8Z4B, and lower-resolution 4GBK. No significant outliers are observed in disallowed regions, indicating that local stereochemical quality is preserved under both ambient and cryogenic conditions. Background contour maps indicate energetically favored and allowed regions of Ramachandran space, and observed ϕ/ψ values are shown as scatter points. Across all datasets, the large majority of residues occupy favored regions, and expected residue-specific distribution patterns are retained.

Collectively, the comparative Ramachandran analyses of detemir and aspart insulins clearly demonstrate that both analogs consistently preserve their local backbone geometries under different temperature and data-collection regimes. The relative broadening of angular clusters from cryogenic to ambient-temperature datasets, consistent with increased thermal mobility yet remaining within accepted energetic limits, appears to represent a common dynamic feature in both molecules [34]. The most striking molecule-specific difference between the analyses might be the complete absence of residues (n=0) in the aspart contour maps, reflecting the B28-Proline → Aspartic acid mutation in aspart insulin [35], in contrast to the proline distribution (n=4) observed in detemir. It is highly noteworthy that the macroscopic lattice disorders, which manifest as pseudo-translational symmetry (tNCS) for detemir and severe merohedral twinning for aspart in the reciprocal-space analyses [26], do not lead to any stereochemical deviations (outliers) in the Ramachandran plots [33]. This suggests that the strong lattice heterogeneity observed via multicrystal approaches at near-physiological ambient temperatures does not originate from protein misfolding, but rather from the natural structural plasticity [6], [36] of the molecule while maintaining its stable, intrinsic three-dimensional folding.

### 1.7. Analog dependent coupling of lattice-correlated disorder and solvent accessibility

To investigate structural dynamics and intra-lattice motions of detemir insulin under different temperature regimes, two-dimensional diffuse-scattering histograms, one-dimensional logarithmic intensity profiles, and atom-level solvent-accessible surface-area plots were generated (**Fig. 5**). In contrast to Bragg reflections, which report the average electron-density distribution in the crystal, X-ray diffuse scattering is sensitive to spatially correlated electron-density variations and therefore provides information on molecular flexibility and lattice disorder [37], [38], [39].

**Figure 5.**
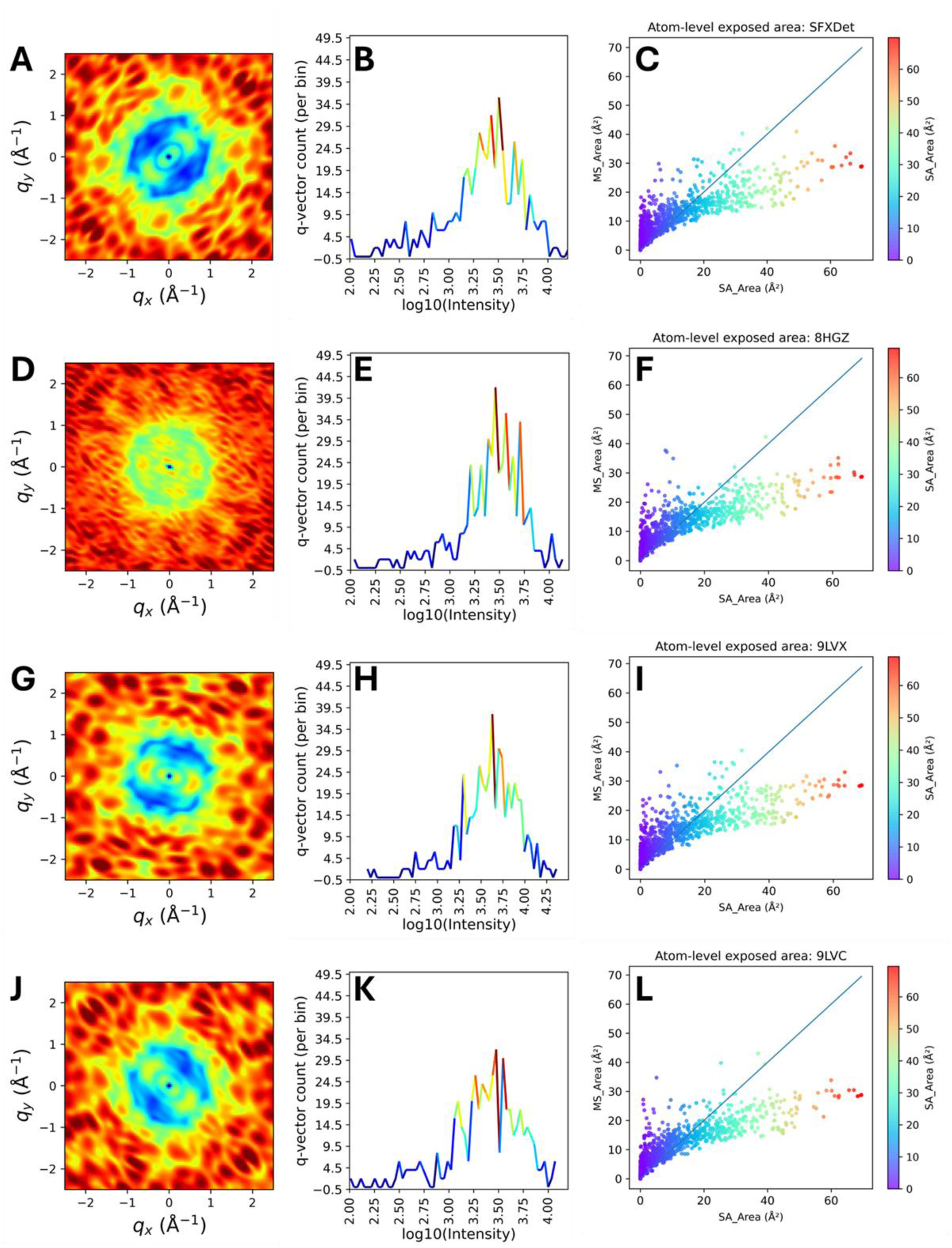
Comparative diffuse-scattering and atom-level surface-accessibility analysis of detemir insulin under ambient-temperature and cryogenic conditions. Panels A–C show ambient-temperature SFX: two-dimensional diffuse-scattering map (A), one-dimensional logarithmic intensity profile (B), and atom-level exposed-area relationship plot (C). Panels D–F show cryogenic single-crystal 8HGZ. Panels G–I show ambient-temperature multicrystal 9LVX. Panels J–L show cryogenic single-crystal 9LVC. In two-dimensional diffuse maps (A, D, G, J), color ranges from dark blue (low intensity) to red (high intensity). Ambient-temperature datasets (A, G) display more continuous and anisotropic diffuse features, consistent with stronger correlated motions, whereas cryogenic datasets (D, J) show a more restricted diffuse signal. One-dimensional logarithmic intensity profiles (B, E, H, K) summarize diffuse-intensity distributions in reciprocal space [log_10_(Intensity) versus *q*-vector count]. Ambient-temperature profiles (B, H) are smoother and more continuous, whereas cryogenic profiles (E, K) are more discrete, consistent with reduced conformational sampling after cryogenic trapping. Atom-level plots (C, F, I, L) compare solvent-accessible surface area (*SA_Area*) with molecular surface area (*MS_Area*). Cryogenic datasets (F, L) show tighter clustering, whereas ambient-temperature datasets (C, I), particularly SFX, show relatively broader dispersion, consistent with greater conformational plasticity and local breathing motions at near-physiological temperature.

Inspection of the two-dimensional diffuse maps (**Fig. 5A, 5D, 5G, 5J**) together with the corresponding one-dimensional distributions reveals a clear dynamic contrast between ambient-temperature datasets (SFX, 9LVX) and cryogenic datasets (8HGZ, 9LVC). The ambient-temperature maps (**Fig. 5A and 5G**), in which the crystal is not cryo-trapped, show more continuous and anisotropic diffuse features, consistent with increased thermal motion and intra-protein structural fluctuations [40]. By contrast, cryogenic datasets 8HGZ (**Fig. 5D**) and 9LVC (**Fig. 5J**) show a more restricted diffuse profile, consistent with partial suppression of conformational dynamics during cryogenic trapping. These observations suggest that ambient-temperature SFX is better positioned to capture near-physiological thermal fluctuations and correlated atomic displacements within the detemir lattice.

This map-level contrast is further supported by comparison of one-dimensional logarithmic intensity histograms [*log*_10_(*I*)] as a function of *q*-vector count (**Fig. 5B, 5E, 5H, 5K**). These distributions indicate how frequently a given diffuse intensity occurs across reciprocal space. For ambient-temperature datasets (**Fig. 5B and Fig. 5H**), the profiles are smoother and more continuous, consistent with sampling of a broader and more continuous dynamic energy landscape. In cryogenic datasets (**Fig. 5E and Fig. 5K**), profiles appear more discrete, with sharper peak structure, consistent with preferential trapping of lower-energy substates when thermal motion is reduced [9], [41]. The color schemes in **Fig. 5** were selected to emphasize these crystallographic features. In two-dimensional diffuse maps (**Fig. 5A, 5D, 5G, 5J**), color ranges from low intensity (dark blue) to high intensity (red), with higher-intensity regions likely corresponding to stronger correlated motions and local lattice flexibility. In one-dimensional histograms (**Fig. 5B, 5E, 5H, 5K**), line color is linked to vector count (jet scale): red peak regions indicate intensities represented by a high number of *q* vectors, whereas blue trough regions indicate comparatively rare intensity levels.

To examine whether correlated mobility inferred from diffuse scattering is reflected in protein–solvent exposure, scatter plots of solvent-accessible surface area (SA_Area_) versus molecular surface area (MS_Area_) were analyzed at the atom level (**Fig. 5C, 5F, 5I, 5L**) [42]. Cryogenic single-crystal datasets (**Fig. 5F** and **5L**) show relatively tighter clustering around a narrower trend, consistent with thermal contraction and increased lattice compaction during freezing. In contrast, ambient-temperature SFX (**Fig. 5C**) and multicrystal 9LVX (**Fig. 5I**) show relative broader dispersion, consistent with increased thermal flexibility. The wider SA/MS distribution, particularly in SFX, is compatible with local conformational “breathing” in detemir hexamers and potentially increased transient surface exposure.

Collectively, these results indicate an association between stronger diffuse-scattering features at ambient temperature and broader atom-level surface accessibility. Whereas cryogenic structures may preferentially represent lower-energy, more static substates, ambient-temperature approaches such as SFX appear to capture a wider range of intrinsic detemir dynamics that may be relevant to its protracted pharmacological behavior and dissociation equilibrium [43].

Likewise, to evaluate differences in model-derived diffuse-scattering behavior across aspart insulin datasets, **Figure 6** compares ambient-temperature SFX (panels **A–C**), cryogenic single-crystal 4GBK (panels **D–F**) [6], cryogenic single-crystal 4GBC (panels **G–I**) [6], and ambient-temperature multicrystal 8Z4B (panels **J–L**) [36]. The figure includes two-dimensional diffuse maps (panels **A, D, G, J**), one-dimensional log-intensity histograms (panels **B, E, H, K**), and atom-level SA_Area_ versus MS_Area_ scatter plots (panels **C, F, I, L**). The diffuse intensity for **Figures 5** and **6** is computed using a liquid-like-motion/Gaussian formalism [44], as described in the method section. Accordingly, panels **A, D, G,** and **J** should be interpreted as model-derived diffuse representations rather than direct detector-level diffuse-scattering measurements. Within this framework, all four datasets show anisotropic diffuse features, with visible differences in cloud morphology and intensity texture between conditions. Ambient-temperature SFX (**Fig. 6A**) and ambient-temperature multicrystal 8Z4B (**Fig. 6J**) show relatively broader diffuse clouds than cryogenic 4GBK (**Fig. 6D**) and 4GBC (**Fig. 6G**), whereas cryogenic maps appear spatially restricted. Panels **B, E, H,** and **K** are histograms of positive diffuse intensities *log*_10_(*I*) plotted against per-bin *q*-vector counts as previously mentioned. These profiles therefore represent the statistical distribution of modeled diffuse intensities, not direct intensity-versus- *q* traces. Cryogenic datasets (panels **E, H**) display sharper, more localized peaks, whereas ambient-temperature datasets (panels **B, K**) show broader, multimodal profiles, consistent with wider sampling of the intensity distribution in the model space.

**Figure 6.**
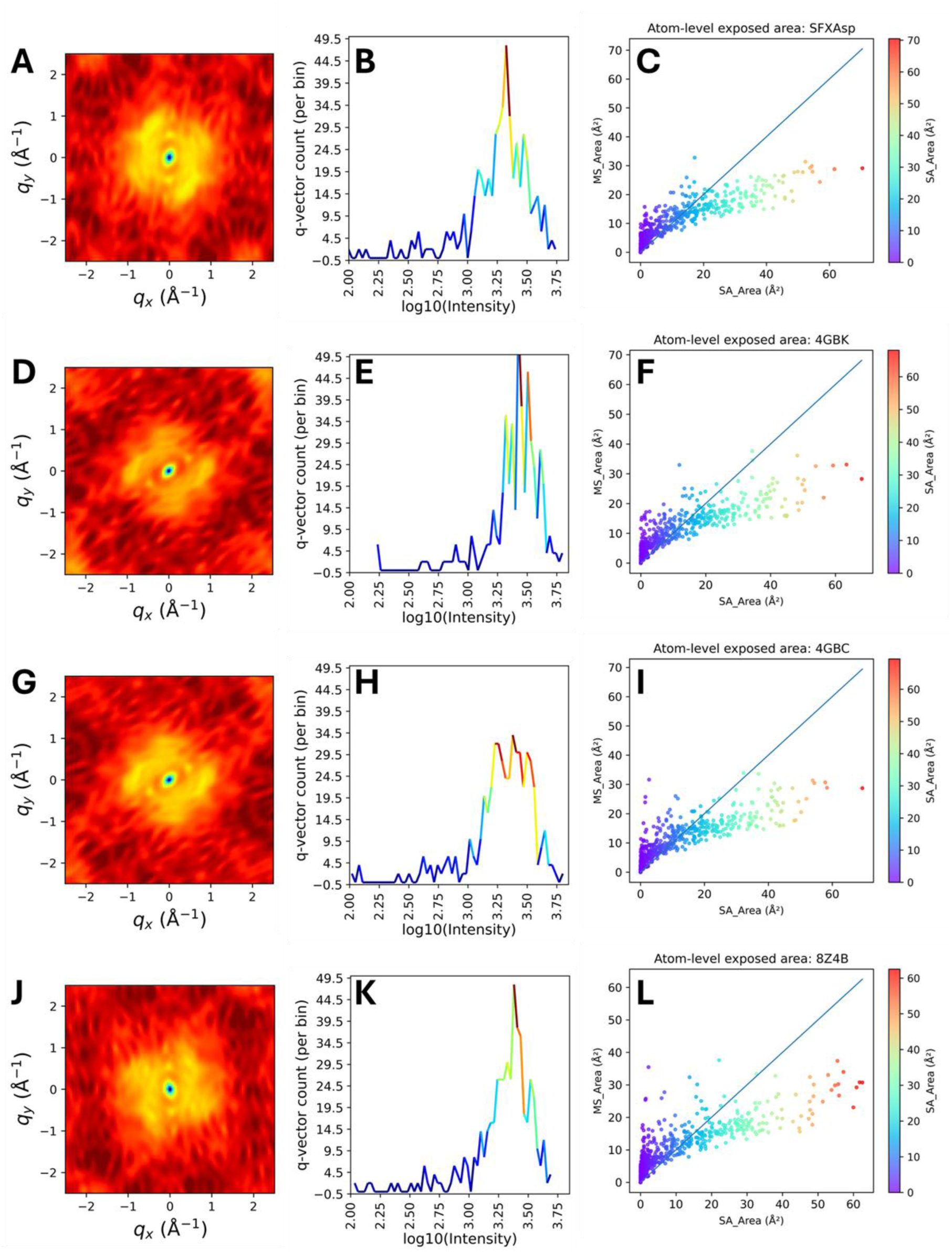
Comparative model-derived diffuse-scattering and atom-level surface-accessibility analysis of aspart insulin datasets collected under ambient-temperature and cryogenic conditions. Panels A–C: ambient-temperature SFX (new dataset); panels D–F and panels G–I: cryogenic single-crystal 4GBK and 4GBC; panels J–L: ambient-temperature multicrystal 8Z4B. For each dataset, the left panel shows the two-dimensional diffuse map (*q_x_ q*_y_); the middle panel shows the histogram of positive diffuse intensities as log_10_(I) versus per-bin q-vector count; and the right panel shows atom-level SA_Area_ versus MS_Area_ with the y=x reference line. Diffuse maps are generated using the liquid-like-motion/Gaussian framework and therefore represent model-derived reciprocal-space features rather than direct detector diffuse-scattering measurements. Across datasets, diffuse-map anisotropy, intensity-distribution shape, and the dispersion of SA versus MS differ systematically, indicating condition-dependent differences in inferred lattice-correlated disorder and surface-accessibility heterogeneity.

Atom-level SA_Area_– MS_Area_ plots (panels **C, F, I, L**) show positive SA/MS association in all datasets [45]. In the present panel set, SFX (**Fig. 6C**) is comparatively tighter around the central trend, whereas 4GBK (**Fig. 6F**), 4GBC (**Fig. 6I**), and 8Z4B (**Fig. 6L**) show broader downward spread from the *y* = *x* reference line at larger SA_Area_ values.

Taken together, **Figure 6** indicates dataset-dependent differences in model-derived diffuse-scattering statistics and atom-level surface-area dispersion across the four aspart structures. These differences are quantitatively observable in the generated representations; however, assignment to specific mechanistic substates should be made cautiously unless supported by additional experimental diffuse-scattering validation and independent ensemble modeling [46].

Collectively, our comparative analysis of the detemir (**Fig. 5**) and aspart (**Fig. 6**) datasets indicates a shared temperature-dependent pattern in model-derived diffuse-scattering behavior, with ambient-temperature datasets in both analogs showing modest but consistent differences than their cryogenic counterparts [44], [47]. This overall trend is consistent with enhanced conformational sampling under non-cryo-trapped conditions and reduced motional heterogeneity after cryogenic trapping [47]. At the same time, the atom-level SA_Area_ – MS_Area_ relationships [45] reveal a notable analog-specific distinction: whereas ambient-temperature detemir datasets (particularly SFX; **Fig. 5C**) show broader surface-area dispersion relative to cryogenic datasets (**Fig. 5F, 5L**), aspart SFX (**Fig. 6C**) appears comparatively tighter around the central SA/MS trend, with broader dispersion more evident in 4GBK, 4GBC, and 8Z4B (**Fig. 6F, 6I, 6L**). Thus, while both systems exhibit consistent ambient-versus-cryogenic contrast in diffuse-scattering statistics, their solvent-exposure distributions are not identical, suggesting that the coupling between lattice-correlated disorder and local surface accessibility might be molecule dependent [48]. Given that the diffuse maps are generated within a liquid-like-motion/Gaussian framework [40], [44], these observations are most appropriately interpreted as robust comparative signatures at the level of model-derived covariance structure [44], pending orthogonal validation by experimental diffuse-scattering and ensemble-based refinement [30], [49].

To bridge these lattice-wide covariance signatures with a site-specific metric, we next applied RABDAM analysis [50]. Although RABDAM was originally developed to identify specific radiation-damage hotspots, its BDamage formalism normalizes atomic B-factors by local packing environment and is therefore sensitive to localized flexibility outliers. For detemir, mean BDamage values are near unity across datasets (SFX 1.0025; 8HGZ 0.9996; 9LVX 0.9997; 9LVC 0.9995), while maximum values vary strongly (SFX 3.6587; 8HGZ 2.9651; 9LVX 2.1086; 9LVC 2.4384), indicating markedly stronger localized extremes in the ambient-temperature SFX model. For aspart, mean BDamage values are likewise near unity (SFX 0.9995; 4GBK 0.9989; 4GBC 0.9998; 8Z4B 0.9991), but maximum values show a different ranking (SFX 1.6346; 4GBK 2.1679; 4GBC 1.6278; 8Z4B 3.2295), with the highest localized extremes in 8Z4B and comparatively lower maxima in SFX and 4GBC. Collectively, these data support a conservative interpretation: global normalized BDamage behavior is similar across each analog, whereas the magnitude of site-specific high-BDamage outliers is dataset-dependent. In practical terms, these analog-specific outlier patterns provide residue-level targets for stability monitoring and formulation optimization, and they might help link ambient-versus-cryogenic heterogeneity to clinically relevant behavior such as protracted detemir assembly versus faster-acting aspart dynamics.

## 2. Discussion

Insulin detemir and insulin aspart are widely regarded as pharmacologically contrasting kinetic designs: detemir is optimized for prolonged basal activity through acylation-dependent self-association and albumin binding [9], [51], whereas aspart is optimized for rapid prandial action through reduced self-association and faster monomer availability [6], [52]. In this comparative crystallographic framework, these design principles are *reflected* in different structural/disorder signatures rather than interpreted as direct one-to-one causal determinants of in vivo kinetics.

Across both analogs, comparison of ambient-temperature SFX/multicrystal datasets with cryogenic single-crystal PDB structures shows a shared temperature-dependent trend: expanded unit-cell dimensions (**Table 1, Table S1-S3**), broader reciprocal-space heterogeneity (**Fig. 1** and **Fig. 2**), and broader/more anisotropic model-derived diffuse patterns (**Fig. 5** and **Fig. 6**) under ambient conditions. Consistent with prior room-temperature crystallography literature, these features support increased conformational sampling relative to cryogenic trapping. At the same time, Ramachandran distributions remain within acceptable stereochemical limits, indicating that increased disorder signatures are not attributable to gross misfolding of backbone geometry (**Fig. 3** and **Fig. 4**). Despite this common ambient-versus-cryogenic trend, analog-specific differences are evident. For aspart, ambient-condition datasets show a strong propensity for merohedral twinning (in several cases approaching the twinned limit), consistent with lattice-level domain intergrowth in the trigonal setting [6], [25], [27]. For detemir, disorder is more prominently expressed through pseudo-translational modulation (tNCS) together with partial twinning, consistent with its distinct dihexamer-associated packing context [53], [54], [55]. Thus, the dominant crystallographic “carrier” of disorder differs between analogs, even when both are measured under near-physiological conditions. Divergence is also observed at the atomistic surface/disorder level. In the SA–MS analyses (**Fig. 5** and **Fig. 6**), ambient detemir SFX exhibits broader surface-accessibility dispersion (**Fig. 5C**), whereas aspart SFX appears comparatively tighter around the central trend (**Fig. 6C**), with broader spread more evident in other aspart datasets (**Fig. 6F, 6I, 6L**). RABDAM is consistent with this distinction: although mean BDamage values remain near unity across datasets (indicating comparable global normalization), maximum BDamage outliers differ by analog and condition, with particularly strong localized extremes in detemir SFX and a different ranking pattern in aspart. These findings support a conservative interpretation that global disorder baselines are comparable, but the magnitude and distribution of localized vulnerable/flexible sites are dataset- and molecule-dependent. Under SFX specifically, both analogs share ensemble-like signatures relative to legacy cryogenic structures (greater heterogeneity, broader diffuse-derived distributions, and stronger sensitivity to twinning/disorder metrics), but they diverge in how disorder is partitioned between lattice-level effects (tNCS versus twinning-dominant behavior) and local accessibility/mobility outliers. This shared-plus-divergent pattern is important because it links ambient crystallographic observables to analog-specific structural landscapes without overextending to unsupported mechanistic claims.

Collectively, a biotechnological and medical perspective of the current results propose that ambient/SFX structural characterization might provide practical readouts for insulin development beyond static cryogenic models (**Fig. 7**). Read as a structural timeline, detemir evolves from the foundational cryogenic framework of 1XDA through high-resolution ordered references (8HGZ, 9LVC) to the more heterogeneous ambient states (9LVX) and finally to near-physiological ensemble capture by SFX, whereas aspart progresses from cryogenic 4GBK/4GBC reference states to ambient multicrystal 8Z4B and SFX representations that emphasize condition-dependent disorder partitioning. This longitudinal PDB-to-SFX view links crystallographic descriptors to analog-specific functional design: protracted behavior in detemir might be associated with packing-and disorder-signature profiles distinct from the rapid-acting aspart landscape. Quantitative signatures such as twinning tendency, pseudo-translation metrics, model-derived diffuse anisotropy, SA–MS dispersion, and BDamage outlier profiles therefore provide useful comparators for formulation screening, stability assessment, and engineering of analog-specific assembly behavior. Namely, the study supports a shift from a single-structure paradigm toward a function-relevant ensemble perspective for the design of next-generation rapid-acting and protracted insulins.

**Figure 7.**
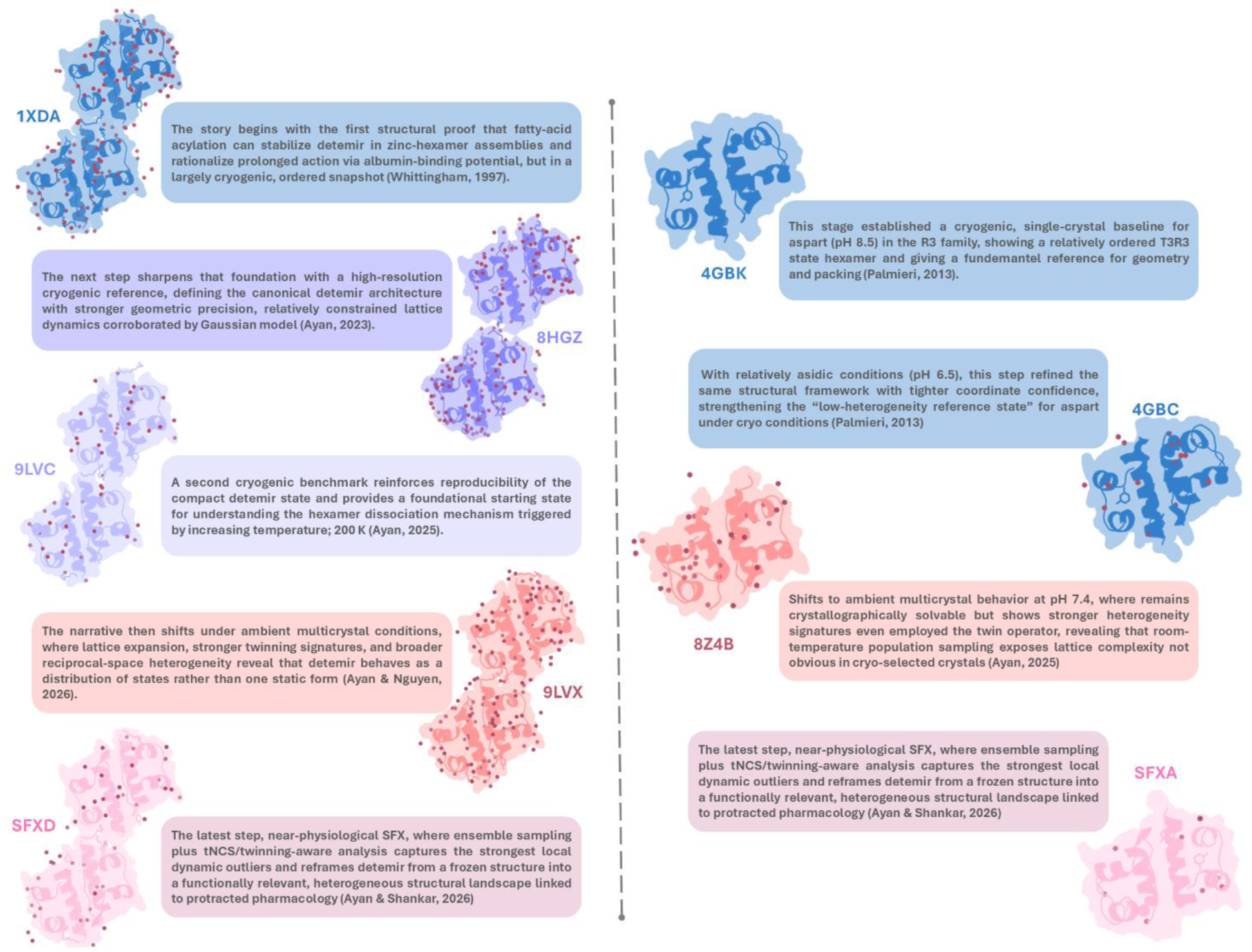
Evolutionary structural timeline of insulin analog crystallography from cryogenic single-crystal references to ambient/near-physiological ensemble methods. Left panel (detemir): progression from 1XDA to 8HGZ, 9LVC, 9LVX, and SFXD, highlighting the shift from compact cryogenic reference states to ambient multicrystal/SFX datasets with greater lattice heterogeneity, twinning sensitivity, and ensemble-level dynamics. Right panel (aspart): progression from 4GBK and 4GBC to 8Z4B and SFXA, showing a parallel transition from ordered cryogenic type reference structures to ambient-condition datasets that better capture heterogeneity. Together, the two trajectories illustrate a methodological and conceptual transition from static, low-temperature snapshots to physiologically relevant structural ensembles for interpreting protracted (detemir) versus rapid-acting (aspart) behavior.

## 3. Method

### 3.1. Sample Preparation and Crystallization

NovoRapid® (insulin aspart) and Levemir® (insulin detemir) were crystallized using a sitting-drop microbatch-under-oil vapor-diffusion approach, following the workflow reported by Ayan et al. [56], [57], [58]. In brief, each protein sample was combined with approximately 3,500 sparse-matrix screening conditions at a 1:1 (v/v) ratio at room temperature, then dispensed into 72-well Terasaki plates (Greiner-Bio, Germany). Each 0.83 µL drop was covered with 16.6 µL paraffin oil (Tekkim Kimya, Türkiye) and incubated at ambient conditions. Crystal growth was tracked by light microscopy, and well-formed crystals were typically observed within 48 h. The best crystals were obtained from a mother liquor containing 0.2 M sodium acetate trihydrate and 0.1 M Tris-HCl (pH 8.5). Additional trays were prepared under the same screening setup to ensure reproducibility. After optimizing a seeding strategy for oil-under-microbatch crystals, crystallization was scaled to a 1 mL batch volume, where microcrystal formation was verified by compound light microscopy.

### 3.2. Sample delivery, data collection, and processing

To obtain the crystal density required for serial injection (approximately 10^8^ crystals per injection), a 1 mL microcrystal slurry aliquot was centrifuged at 3,000 rpm for 5 min. After centrifugation, 990 µL of supernatant was removed, leaving a 10 µL pellet. This concentrated fraction was mixed with Synco Chemical grease (No. 42150) on a plastic plate. The grease–crystal mixture was then gently transferred into a 200 µL sample cartridge with a spatula, taking care to avoid bubble formation. The cartridge was installed on a high-vacuum injector fitted with a 75 µm inner-diameter nozzle and operated inside the helium-filled DAPHNIS diffraction chamber [59]. Diffraction data for insulin detemir and insulin aspart crystals were collected at SACLA BL2 Experimental Hutch 3 using 10 keV XFEL pulses (<10 fs, 30 Hz), with the MPCCD detector positioned 50 mm from the sample. Buffer was supplied by HPLC at 0.0127 µL/min, while the grease-based crystal suspension was extruded at 0.79 µL/min. Raw frames were screened for crystal hits with `peakfinder8` in *Cheetah* [60], [61]. Accepted hits were indexed in *CrystFEL* v0.9.1 [62] using `xgandalf` (min-snr = 4.5, threshold = 100, minimum pixel count = 2). Indexed patterns were subsequently scaled and merged with partialator (`--model=unity`) in *CrystFEL* to obtain structure-factor amplitudes. Data-quality statistics were evaluated using `comparehkl` and `checkhkl` (**Table S1-S2**). For both detemir and aspart datasets, the indexing ambiguity identified by `cell_tool` was resolved by applying the ambigator operator `-h,-h-k,-l`. Structures were determined by automated molecular replacement in *Phaser*, yielding space group R3:H for both detemir and aspart. Initial rigid-body refinement employed previously reported cryogenic models (PDB ID: 1XDA for detemir; PDB ID: 4GBK for aspart R3:H). After simulated annealing refinement, the individual atomic coordinates and TLS parameters were further optimized. Solvent models were curated manually in *Coot* by retaining waters supported by clear difference density and removing unsupported sites. Data-collection and refinement statistics are summarized in Tables 1–3, and structural figures were prepared in *PyMOL*.

### 3.3. Reciprocal space mapping and intensity field reconstruction

To visualize lattice heterogeneity, reflection broadening, and reciprocal-space sampling variations across the datasets, reciprocal-space mapping was performed using our custom-developed Python script, which analyzed Bragg data (extracted from MTZ files) and model-derived calculated structure factors (*F*_*calc*_) [28]. In three-dimensional reciprocal space, the positional coordinate of each reflection is defined by its scattering vector ***q***. For a given set of Miller indices (*h*, *k*, *l*), the scattering vector is calculated using the reciprocal lattice basis vectors (*a* ∗, *b* ∗, *c* ∗)according to the following equation:

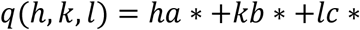

The scattering intensity *I*(*q*) at each reciprocal lattice point is directly proportional to the square of the structure factor amplitude (rec10):

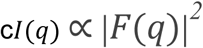

Conventional two-dimensional and three-dimensional reciprocal-space mapping relies on projecting the integrated scattering intensity onto a specific plane or volume to extract structural details that are often obscured in standard profile analyses [28]. Instead, to construct continuous two-dimensional section plots (e.g., *HK*0, *H*0*L*, 0*KL*) and three-dimensional reciprocal lattice point clouds, we applied a Gaussian kernel broadening algorithm to the discrete Bragg nodes [63]. This deterministic broadening (using a Gaussian sigma of 1.8 pixels on a 350-bin 2D grid) spreads the discrete points into a continuous, regularized reciprocal-intensity field. Using this framework, kernel-broadened observed fields (*I*_*obs*_) and calculated regularized reciprocal-space intensity representations derived from the model (*F*_*calc*_) were generated. To quantitatively assess the deviations between the experimental data and the model, linear difference maps (*I*_*obs*_ − *I*_*calc*_) and logarithmic ratio maps (*log*_10_(*I*_*obs*_/*I*_*calc*_) were computed. It is important to note that these mathematically reconstructed maps represent the statistical distribution and volumetric shape of modeled reciprocal-space intensities [30]. Therefore, they serve as regularized representations of macroscopic lattice disorder and reciprocal-space heterogeneity, rather than direct instrumental measurements of diffuse scattering [28].

### 3.4. Ramachandran plot analysis and stereochemical validation

To evaluate the stereochemical quality and local backbone geometry of the refined models, Ramachandran plot analysis was conducted using our custom-developed Python script. The primary main-chain dihedral angles, *ϕ*(*C*_*i*−1_ − *N* − *C*_*⍺*_ − *C*_*i*_), and *ψ*(*N* − *C*_*⍺*_ − *C* − *N*_*i*+1_) were extracted for each residue and rigorously filtered to ensure all data points fell within the valid [−180°, 180°] torsional boundaries [34]. To visually and quantitatively map the conformational space against expected steric limits, a continuous pseudo-energy background was computationally generated. Inspired by classical hard-sphere models and energetic distributions of allowed backbone conformations [33], [34], the sterically favored and partially allowed regions were mathematically approximated using a superposition of two-dimensional Gaussian functions evaluated over a continuous 200 ✕ 200 grid. The pseudo-energy landscape, *Z*(*ϕ*, *ψ*), was calculated according to the following formulated equation:

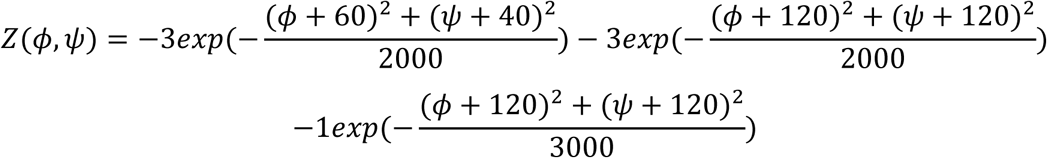

This generated scalar field was subsequently min-max normalized to a scale, establishing a gradient contour map that distinguishes the highly populated core (energetically favored) regions from the partially allowed and sterically restricted (disallowed) zones [33]. Recognizing that different amino acid side chains impose distinct steric hindrances on the polypeptide backbone, the structural analysis was stratified by residue type [34]. Specifically, glycine residues, which lack a chiral side chain and therefore possess significantly greater conformational freedom, along with proline residues, whose cyclic pyrrolidine ring severely constrains the *ϕ* angle, were isolated and plotted independently from the general amino acid population. Finally, individual residues were categorized and color-coded as ‘Favored’, ‘Allowed’, or ‘Outlier’ according to standard stereochemical criteria [33], enabling a robust assessment of backbone integrity and local folding anomalies across the respective datasets.

## Conclusion

This study establishes a comparative framework for evaluating insulin analog disorder across near-physiological and cryogenic crystallographic regimes. Detemir and aspart share a robust ambient-condition signature of expanded heterogeneity, but they diverge in dominant disorder mechanisms: detemir is characterized by pronounced pseudo-translational modulation with moderate twinning, while aspart is dominated by strong merohedral twinning with minimal tNCS contribution. These distinctions remain visible across reciprocal-space analyses, model-derived diffuse statistics, and residue-level BDamage outlier behavior, while overall stereochemical integrity is preserved in all structures. Methodologically, the work also clarifies an important interpretive boundary: reciprocal-space and diffuse-like maps reconstructed from merged Bragg data are highly informative for comparative heterogeneity, but they should not be conflated with detector-level experimental diffuse scattering. Within this calibrated scope, the integrated metrics used here provide a practical and reproducible panel for cross-dataset benchmarking. Overall, the results advance insulin structural analysis from static single-state interpretation toward a conservative, ensemble-aware perspective more aligned with functional and pharmaceutical reality. This approach is directly relevant for prioritizing analog-specific stability hotspots, refining formulation strategies, and guiding next-generation structure-informed insulin engineering.

## Author Contributions

The insulin analogs were crystallized by E.A. Sample preparation was performed by E.A. under the supervision of J.K. and T.T. SFX data were collected at SACLA XFEL by E.A. and J.K. under the supervision of T.T. and M.Y. Crystal data processing was conducted by E.A. and M.K.S. Structural refinement and interpretation were performed by E.A. Reciprocal space mapping of modeled diffuse scattering was conducted by E.A. under the supervision of M.K.S., using custom scripts developed from the relevant equations reported in the literature, in consultation with the field experts mentioned in the acknowledgments.

## Acknowledgements

This work was supported by the SACLA Research Support Program for Graduate Students (proposals 2023B8058 and 2023B2761) and the Koc University Travel Support Program. We thank Toshinori Yabuuchi, Taito Osaka, Kohei Miyanishi, Gota Yamaguchi, and Tetsuya Ishikawa for support during XFEL data collection, and the Laboratory for Structural Biology at Koc University and the University of Health Sciences for preparing glargine crystalline samples. This study benefited from training received at the International School of Crystallography (ISoC, Erice 2022). E.A. and M.K.S. gratefully acknowledge Michael E. Wall, David Wych, and especially Ariana Peck for valuable assistance in interpreting the diffuse-scattering analyses.

## Data availability

Atomic coordinates and structure-factor data have been deposited in the RCSB Protein Data Bank under accession codes 25HL (detemir) and 25HF (aspart). Additional information is available from the corresponding author upon reasonable request.

## Supplementary Material

**Table S1.** Raw data collection statistics for detemir.

| d_max | d_min | %comp | <l> | <l/sl> | r_mrg | r_meas | r_pim | r_anom | cc1/2 |
| --- | --- | --- | --- | --- | --- | --- | --- | --- | --- |
| 23.55 | 6.11 | 100 | 531.4 | 3 | 0.232 | 0.328 | 0.232 | 0.422 | 0.844 |
| 6.11 | 4.86 | 100 | 603.3 | 3.3 | 0.143 | 0.203 | 0.143 | 0.293 | 0.899 |
| 4.86 | 4.25 | 100 | 1160.5 | 4 | 0.228 | 0.323 | 0.228 | 0.496 | 0.675 |
| 4.25 | 3.87 | 100 | 1043.5 | 3.8 | 0.239 | 0.338 | 0.239 | 0.518 | 0.898 |
| 3.87 | 3.59 | 100 | 792.6 | 3.6 | 0.178 | 0.251 | 0.178 | 0.373 | 0.906 |
| 3.59 | 3.38 | 100 | 681.7 | 3.5 | 0.275 | 0.389 | 0.275 | 0.257 | 0.984 |
| 3.38 | 3.21 | 100 | 495.4 | 3.4 | 0.109 | 0.154 | 0.109 | 0.23 | 0.963 |
| 3.21 | 3.07 | 100 | 370.8 | 3.1 | 0.148 | 0.209 | 0.148 | 0.312 | 0.982 |
| 3.07 | 2.95 | 100 | 318.8 | 3.1 | 0.18 | 0.254 | 0.18 | 0.377 | 0.805 |
| **2.95** | **2.85** | **100** | **261.6** | **3** | **0.209** | **0.296** | **0.209** | **0.477** | **0.665** |
| 23.55 | 2.85 | 100 | 626 | 3.4 | 0.209 | 0.295 | 0.209 | 0.387 | 0.783 |

**Table S2.** Raw data collection statistics for aspart.

| d_max | d_min | %comp | <l> | <l/sl> | r_mrg | r_meas | r_pim | r_anom | cc1/2 |
| --- | --- | --- | --- | --- | --- | --- | --- | --- | --- |
| 23.34 | 4.73 | 100 | 2435.1 | 4.8 | 0.185 | 0.261 | 0.185 | 0.44 | 0.868 |
| 4.73 | 3.76 | 100 | 3906.6 | 5.2 | 0.103 | 0.146 | 0.103 | 0.187 | 0.949 |
| 3.76 | 3.28 | 100 | 2201.9 | 4.5 | 0.164 | 0.231 | 0.164 | 0.363 | 0.758 |
| 3.28 | 2.99 | 100 | 1114.9 | 3.9 | 0.129 | 0.182 | 0.129 | 0.156 | 0.876 |
| 2.99 | 2.77 | 100 | 761.2 | 3.6 | 0.13 | 0.184 | 0.13 | 0.279 | 0.876 |
| 2.77 | 2.61 | 100 | 464.1 | 3.3 | 0.237 | 0.336 | 0.237 | 0.556 | 0.726 |
| 2.61 | 2.48 | 100 | 279 | 2.8 | 0.204 | 0.288 | 0.204 | 0.377 | 0.91 |
| 2.48 | 2.37 | 100 | 172.9 | 2.5 | 0.302 | 0.426 | 0.302 | 0.718 | 0.951 |
| 2.37 | 2.28 | 100 | 128.6 | 2.3 | 0.499 | 0.706 | 0.499 | 1.109 | 0.629 |
| **2.28** | **2.2** | **100** | **84.5** | **1.8** | **0.2** | **0.283** | **0.2** | **0.409** | **0.712** |
| 23.34 | 2.2 | 100 | 1180.5 | 3.5 | 0.151 | 0.214 | 0.151 | 0.29 | 0.916 |

**Table S3.** Data collection and indexing statistics for insulin detemir and aspart.

| sample | hits | hits accepted | indexed | index hit |
| --- | --- | --- | --- | --- |
| detemir | 8932 | 16.1% | 3127 | 28 % |
| aspart | 3823 | 5.7% | 2840 | 74 % |

**Figure S1.**
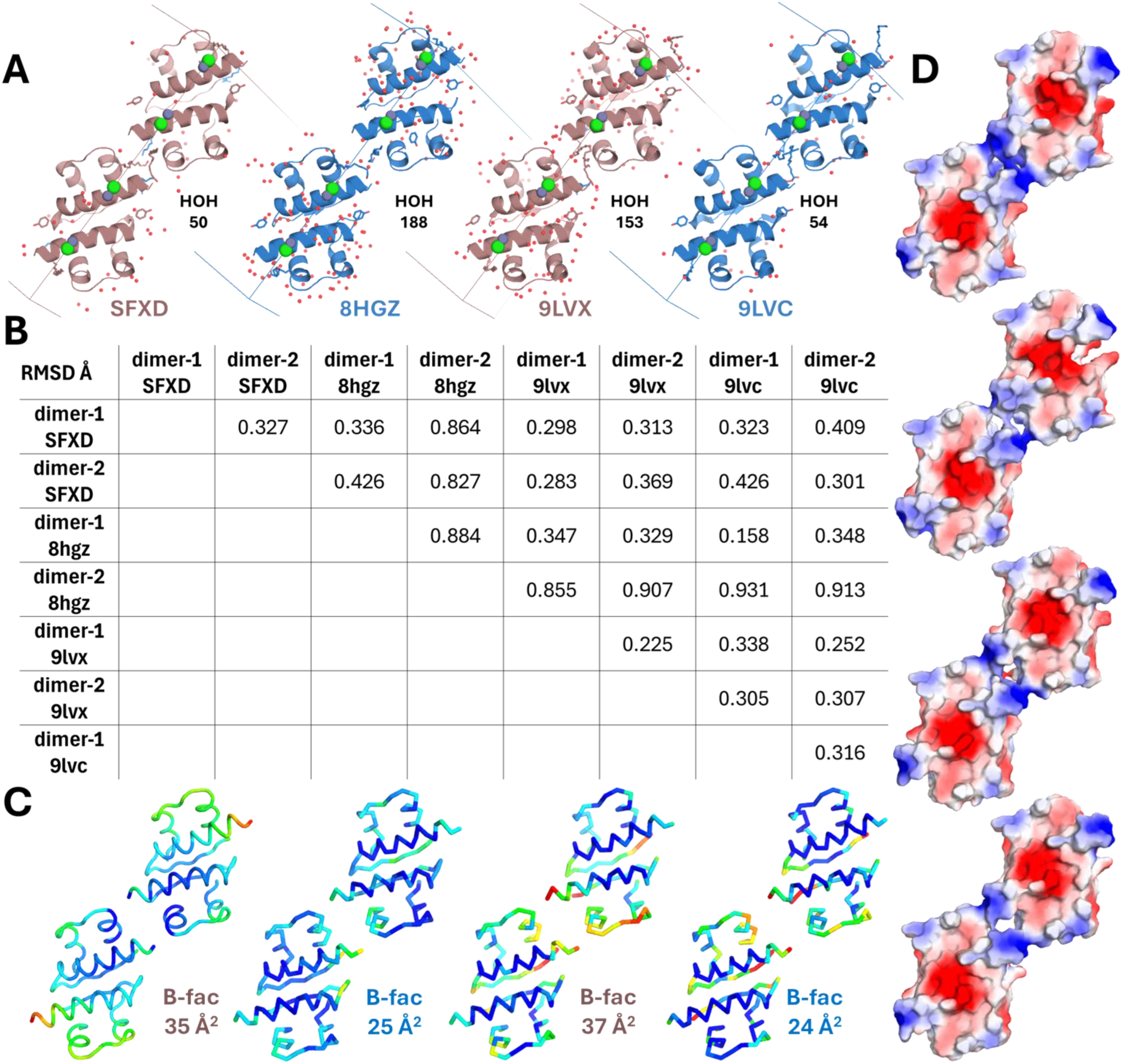
Structural comparison of detemir insulin dimers from ambient-temperature and cryogenic datasets. **(A)** Cartoon representations of the asymmetric-unit dimers from SFXD, 8HGZ, 9LVX, and 9LVC, shown based on the unit cell orientation relative to the three-fold axis. Conserved interfacial water molecules are indicated (red spheres), and representative solvent counts (HOH) are annotated for each model. **(B)** Pairwise *Cα* RMSD matrix (Å) for dimer-1 and dimer-2 from each structure, showing overall close agreement among models and relatively larger deviations between non-equivalent dimer positions. **(C)** Ribbon views colored by atomic displacement parameters (*B*-factors, spectrum range is from 26 to 117), highlighting comparable spatial distributions of flexibility across datasets (mean B values shown for each model). **(D)** Electrostatic surface representations of the corresponding dimers (spectrum range: −65.243 to 65.243), illustrating similar charge-pattern organization at the monomer–monomer interfaces.

**Figure S2.**
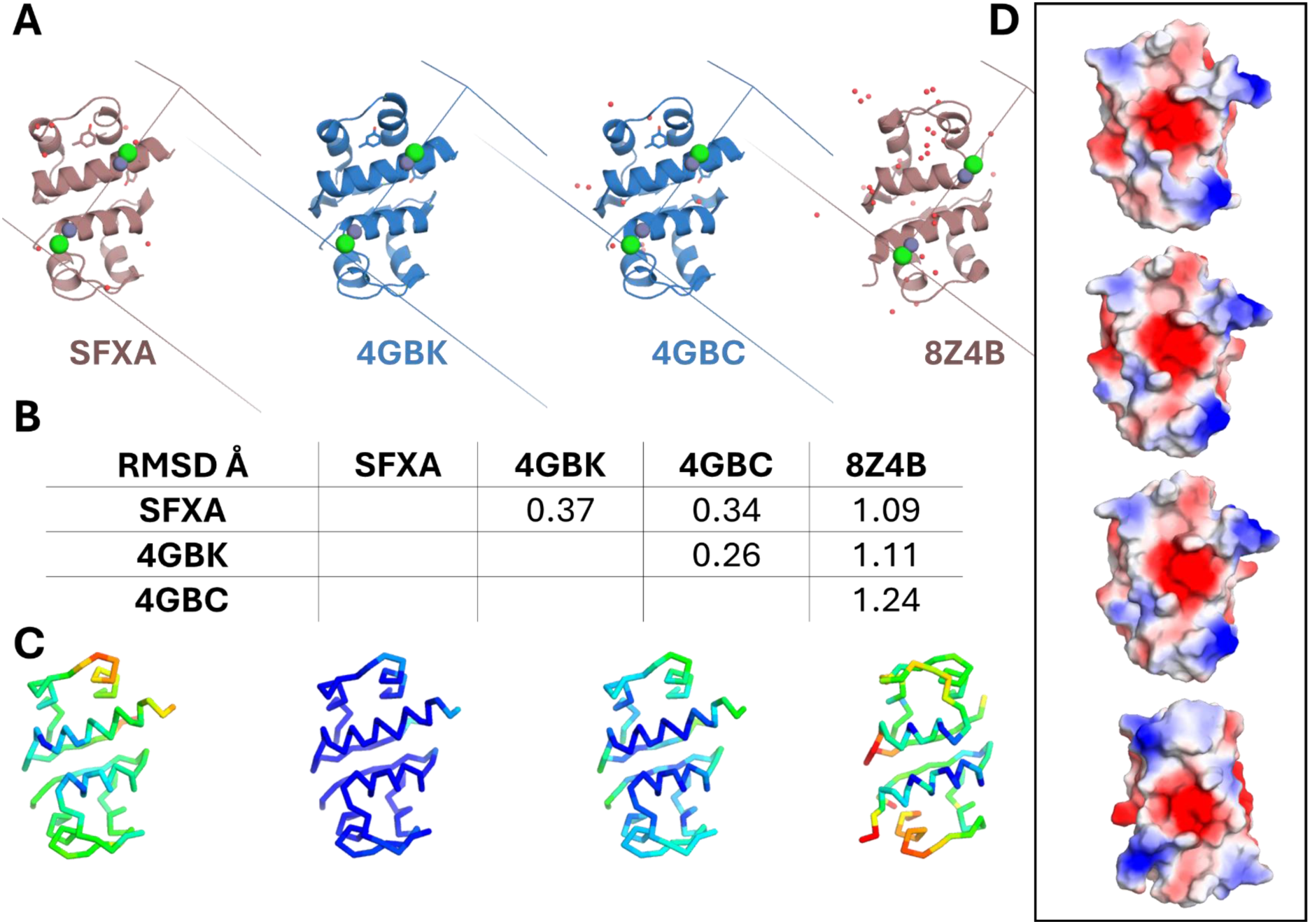
Structural comparison of aspart insulin dimers from ambient-temperature and cryogenic datasets. **(A)** Cartoon representations of the asymmetric-unit dimers from SFXA, 4GBK, 4GBC, and 8Z4B, shown based on the unit cell orientation relative to the three-fold axis. Conserved interfacial water molecules are indicated (red spheres). **(B)** Pairwise *Cα* RMSD matrix (Å) for each structure, showing overall close agreement among models and relatively larger deviations between non-equivalent dimer positions. **(C)** Ribbon views colored by atomic displacement parameters (*B*-factors, spectrum range is from 26 to 117), highlighting comparable spatial distributions of flexibility across datasets (mean B values shown for each model). **(D)** Electrostatic surface representations of the corresponding dimers (spectrum range: −65.243 to 65.243), illustrating similar charge-pattern organization at the monomer–monomer interfaces.

**Figure S3.**
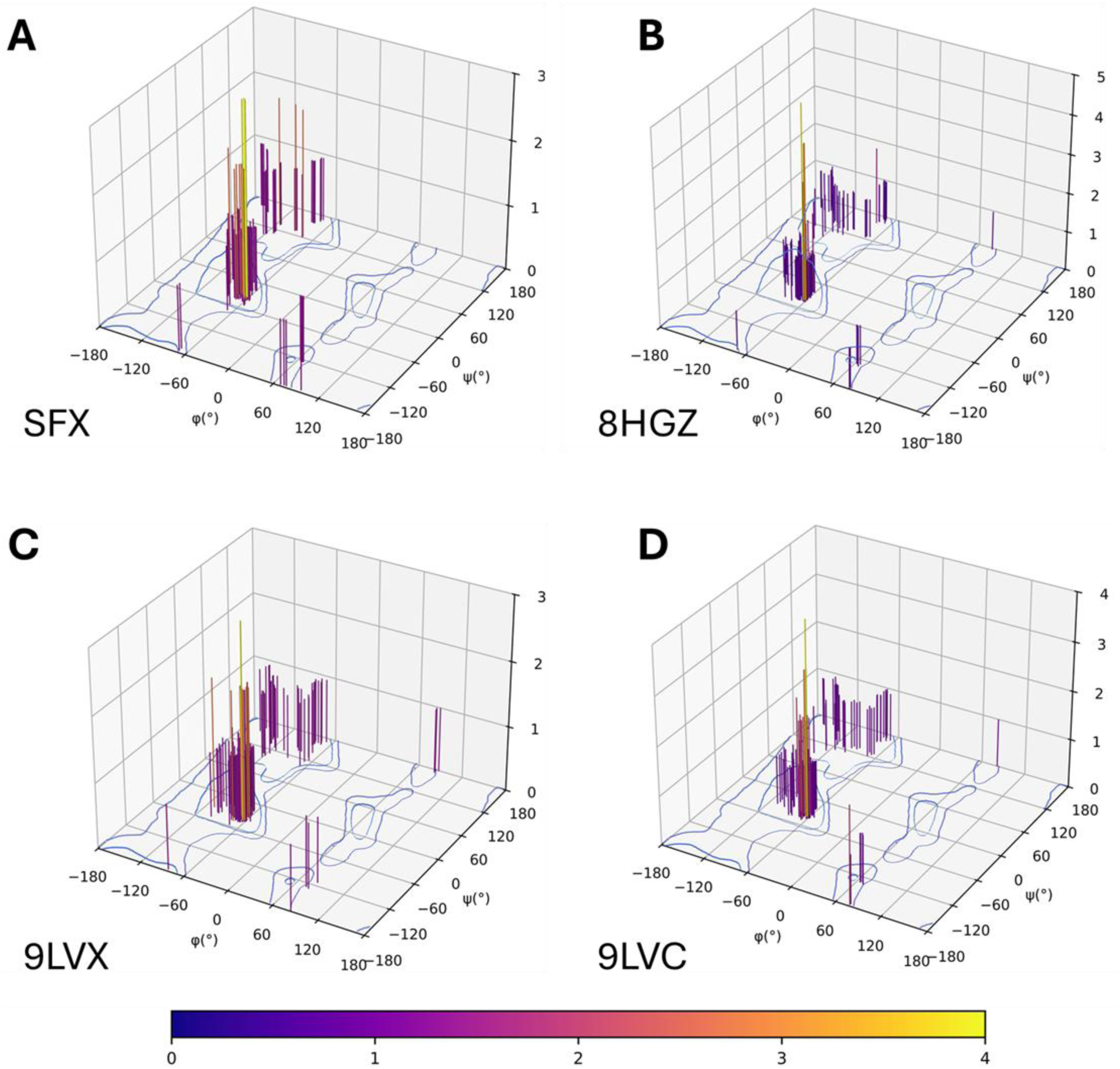
Comparative phi-psi distribution maps for four detemir insulin datasets. **(A)** SFX, **(B)** 8HGZ, **(C)** 9LVX, and **(D)** 9LVC. In each panel, residue-wise observations are plotted in 3D Ramachandran space (ϕ, ψ); vertical bars indicate the corresponding per-point magnitude (same color scale for all panels). The blue contour lines denote reference allowed regions of backbone conformational space. The common plotting scheme enables direct comparison of conformational occupancy and high-value clusters across datasets.

**Figure S4.**
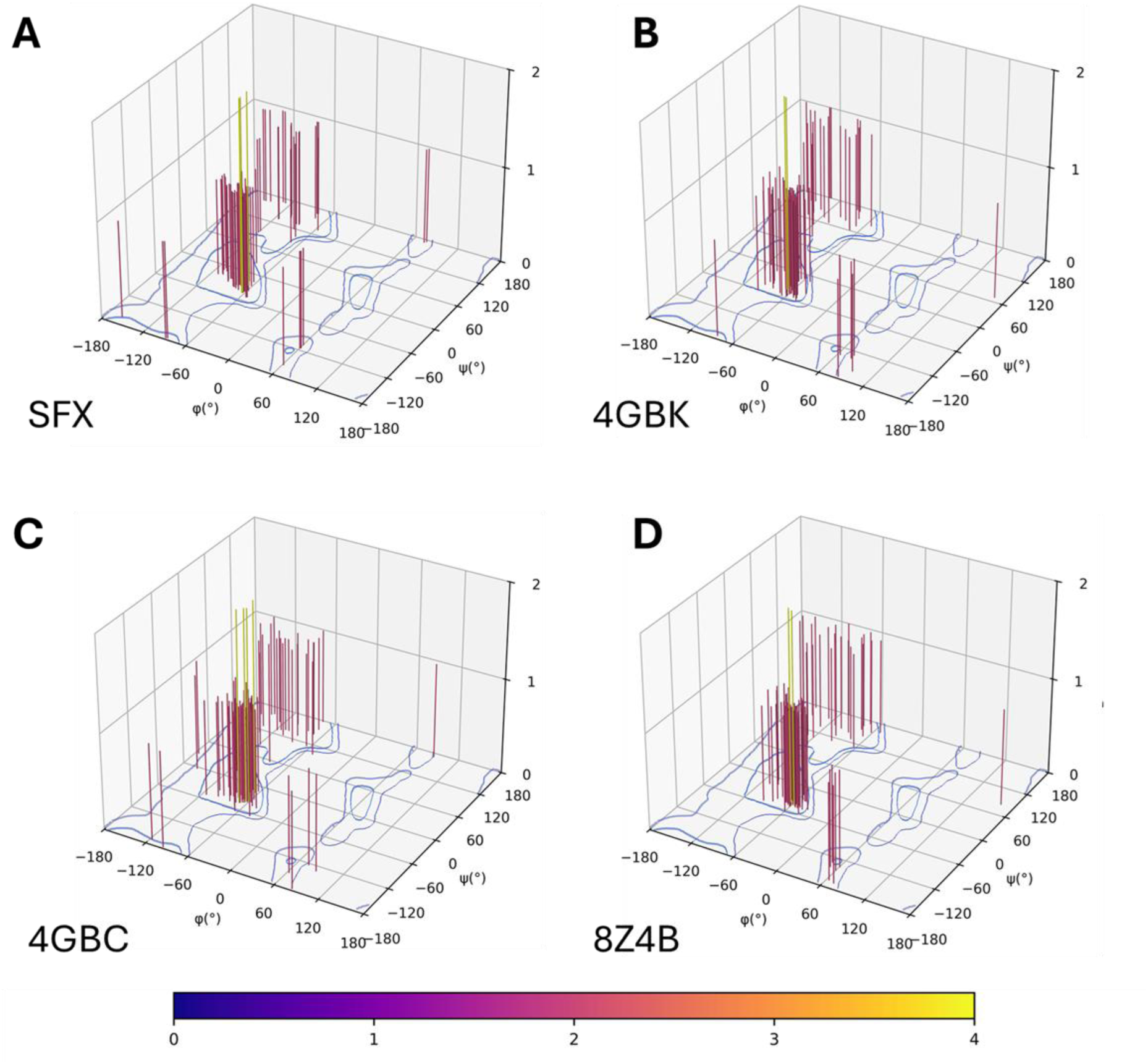
Comparative phi-psi distribution maps for four aspart insulin datasets. **(A)** SFX, **(B)** 8HGZ, **(C)** 9LVX, and **(D)** 9LVC. In each panel, residue-wise observations are plotted in 3D Ramachandran space (ϕ, ψ); vertical bars indicate the corresponding per-point magnitude (same color scale for all panels). The blue contour lines denote reference allowed regions of backbone conformational space. The common plotting scheme enables direct comparison of conformational occupancy and high-value clusters across datasets.

